# A Simulation of Semi-Infectious Particles and Genome Complementation Reproduces Interferon Response by Respiratory Epithelial Cells *in vitro* during Influenza A Virus Infection

**DOI:** 10.64898/2026.05.20.726376

**Authors:** PC Dal-Castel, JD Resnick, JP Sluka, ME Gallagher, M Helfers, IM Bird, JD Ratcliff, SL Grady, JA Glazier

## Abstract

In the respiratory epithelium, interferon (IFN)-induced antiviral resistance acts as a defense against infection. Influenza A viruses (IAVs) have evolved multiple strategies to counteract these defenses, including expression of the viral protein NS1, which inhibits both IFN production and the IFN-mediated transcription of Interferon Stimulated Genes (ISG) in infected cells. However, experiments show that this inhibition is imperfect, especially at a low multiplicity of infection (MOI). One hypothesis to describe this phenomenon relies on the presence of Semi-infectious Particles (SIPs) that fail to express NS1. In this scenario, the IFN response is incompletely suppressed at low MOI, while it is successfully inhibited at high MOI because most cells are infected by multiple virions, allowing complementation to rescue NS1 expression. To test this hypothesis, we developed a computer simulation that models viral gene defects and complementation. We compared the model outputs with *in vitro* experiments at different MOIs. To assess inter-host reproducibility and calibrate the model parameters, we measured IFN levels and viral load over time in bronchial epithelial cell cultures from five human donors. We observed no statistically significant heterogeneity in IFN response or virus production between donors, and the calibrated simulation fits the experimental time series for IFN and viral load. Consistent with literature (1,2), the model predicted higher IFN levels at low MOI than at high MOI. Finally, simulations of IFN treatment applied before and during infection showed reduced viral load, in agreement with our experiments. Increasing the viral genome defect rate above the experimentally estimated rate increased IFN levels and reduced viral load. High MOI simulations showed lower cumulative IFN levels, while NS1 knockout recovered high IFN levels. These results demonstrate the ability of mechanistic models of viral dynamics to predict the innate immune response of epithelial cells during viral infection.

**Author Summary:** Respiratory viruses such as influenza A are highly infectious and pose significant challenges for the human immune system. Through laboratory experiments and computer simulations, we investigated how cells in the respiratory epithelium defend themselves and their neighbors against infection. Using cells collected from different donors, we generated 3-dimensional cell cultures that mimic human airways and measured how they respond to IAV. When a tissue was initially exposed to a small amount of virus, cells could successfully slow or stop the spread of the infection. This phenomenon is hypothesized to be due in part to the high error rate in IAV replication, resulting in many viral particles that are not fully functional. We recapitulated this experimental result with our computational model, validating the model design and parameter estimates. We then simulated a scenario in which cells were pre-treated with interferon, a protective cytokine important to early immune response, and showed that this pre-treatment could successfully limit infection. Laboratory experiments subsequently confirmed this predicted behavior. The computational model reproduced key observations across infection conditions and identified nonfunctional viral particles as important drivers of the early immune response.

## Introduction

Respiratory viruses like Influenza A (IAV) continue to pose a major threat to both human and animal populations, despite the immune response triggered by the cytokines released from the respiratory epithelium early in infection (3,4). Human primary respiratory epithelial cells cultured at the air-liquid-interface (ALI) represent a physiologically relevant model system to study viral infection and host responses, and recapitulate many key features of the airway including diverse cell type composition, polarization, and pseudostratified structure (5,6). In addition to resembling the airway epithelium morphology, these cultures also mimic function, including mucus production, cilia movement, and innate immune responses (5,6). Here, we focus on developing a computer simulation of this *in vitro* model, focusing on IAV-IFN interactions in the epithelium, which are critical to early stages of infection spread and control.

IFN signaling by epithelial cells initiates upon virus internalization (10–12). Toll-like Receptors (TLR) and Retinoic acid-inducible gene I (RIGI) recognize viral double-stranded RNA (dsRNA) and single-stranded RNA (ssRNA), leading to activation of Nuclear Factor-kB (**NF-kB**), and Interferon Regulatory Factors 3 and 7 (**IRF3** and **IRF7**). IRF3 and IRF7 then upregulate **IFN** production, which the infected cells secrete into the extracellular environment (13,14). Neighboring uninfected epithelial cells sense IFN via the Jak-STAT pathway, leading to activation of **IFN-Stimulated Genes (ISG)**. ISGs limit viral spread in the culture by inhibiting viral genome replication, suppressing viral RNA translation, blocking viral entry through endocytosis, and inducing apoptosis of infected cells (15–17). The IAV NS1 protein antagonizes this innate immune response within the cell by inhibiting viral sensing by TLR and RIGI, disrupting IFN signaling via Jak-STAT, suppressing IFN-induced apoptosis, and blocking host gene transcription (2,18–21). We will refer to these viral countermeasures as **antagonism**.

While this antagonism is robust at high **Multiplicity of Infection (MOI)** (22–24), low MOI conditions show significantly elevated IFN levels, with a greater proportion of cells expressing anti-viral genes. One hypothesis for this phenomenon is that NS1 is not 100% effective; however, this does not completely explain why NS1 inhibits IFN production so efficiently at high MOIs (19,20,22,23,25,26). Instead, the incomplete suppression of IFN production in infected cells at low MOI is better explained by a second hypothesis: many infected cells fail to express NS1 due to gene defects in the infecting virions. At high MOI, although these defective virions are still present, most cells are infected by multiple virions, some of which have functional NS1. This allows for complementation: the substitution of defective genes or gene products from one virion by functional genes or gene products from another virion. We will use the term **Infectious Units (IUs)** to refer to complete virions that do not contain any gene defects, and **Noninfectious Particles (NIP)** to refer to virions that contain at least one gene defect. Among NIPs, there are two subcategories. **Semi-Infectious Particles (SIPs)** have one or more damaged or missing segments but can be rescued by coinfection with another SIP or IU that contains a productive version of that gene. **Defective Interfering Particles (DIPs)** are similar in that they have a damaged segment; however, when complemented by another SIP or IU with a functional gene segment, the damaged segment is preferentially replicated, thereby inhibiting successful replication of IUs. In this work, we consider the role of IUs and SIPs, but not the roles of DIPs. See Supplemental Table 1 for more details on biological observations relevant to our work.

The high rate of IAV genome replication errors and segment loss produces abundant and diverse SIPs. In our work, we focus on errors during the replication of the viral genome. Depending on which genes are damaged, these SIPs may fail to carry out critical functions such as genome replication or virus release and can have a large effect on the host’s innate and adaptive immune response after infection. SIPs with missing or defective NS1 are particularly critical to the trajectory of the early innate immune response (27–31). SIPs deficient in NS1 can function as strong inducers of interferon signaling and antiviral resistance in tissues; however, genome complementation with other SIPs or fully infectious particles can restore NS1 suppression of both IFN production and response to extracellular IFN.

Because NS1 targets RNA processing (32), complementation can only rescue inhibition of IFN mRNA production, not translation of host proteins. Therefore, a cell infected by an NS1-competent SIP could produce an IFN response if it produced IFN mRNA before antagonism took place. As such, IFN response varies with the ratio of IUs to the number of cells in the culture at inoculation (i.e., MOI). High MOI infections show more NS1-mediated suppression of IFN production, despite the presence of SIPs, while low MOI infections allow more SIP-driven IFN production at the tissue level. In this sense, including SIPs and viral complementation to a model of IAV infection likely leads to better parametrization and thus better estimates of IFN levels *in vitro* for different MOI.

While experimental manipulation can increase or decrease the frequency of SIPs, they cannot eliminate their production (33). In computer simulations, however, the rate of genome defects leading to SIPs can be freely adjusted, meaning we can predict *in silico* the effect of reducing the generation of SIPs at different MOIs and understand its impact on viral load and IFN levels over time.

Experimental data on IAV infection are fragmented and diverse, spanning multiple species and viral strains (34,35). Building a computational model of IAV infection in nasal epithelial cells *in vitro* using these data requires the systematic integration of this knowledge (36,37). Computational models of micro-scale interactions can clarify how observed macroscale effects emerge. Comparing the effects of individual mechanistic hypotheses in the simulation to *in vitro* experiments can reveal their limitations and guide the design of additional experiments to test these hypotheses. In this work, we consider the following literature-backed hypotheses: 1. SIPs are responsible for high IFN levels (27) – measured as IFN concentration in the supernatant – at low MOI; 2. Complementation rescues inhibition of IFN production at high MOI (2). To test these scenarios, we developed a computer simulation that incorporates SIPs and complementation, calibrated it to experimental time series of IFN levels and viral load, and compared its predictions for high and low MOI conditions and NS1 knockout effects on IFN levels and viral load with in *vitro* results.

Importantly, the computer simulation built in this effort incorporates spatial structure, i.e., the arrangement of cells and/or fields in a coordinate system, where events influence each other with a delay that scales with the distance between them. To justify the importance of spatial structure in IAV infection in epithelial cells in vitro, we first need define the typical distances between infected cells and how long it takes for different signals to travel these distances. The average distance between the nearest initially infected cells at low MOI of 0.01 is of the order of 100μm, or around 10 cell diameters, considering a cell diameter of 10μm (38). Following these estimates, virions released by each infected cell diffuse in a circular region of 50μm in radius on average. Considering a diffusion constant of 6μm^2^/h for virus in mucus (39), the diffusion time for virus to travel a characteristic diffusion length of 50μm is around 100h, as given by *l* = √4 *Dt*, which is larger than the typical 48 hours post-infection (hpi) for viral load peak. If, instead, we consider that viruses spread from cell to cell by contact, and that it takes on average 5h for a cell to initiate release of virions post-infection, the minimum expected travel time is around 25h for the same 50μm radius, which is more consistent with the experimentally observed viral load peak at 48hpi. Due to a faster diffusion coefficient (see Supplemental Table 1), the typical diffusion time for IFNs to travel the same distance is around 2h. If we consider 10h of typical activation time of ISGs in cells exposed to IFN (40), compared to the typical virus travel time, we find that cells further than 3 cell diameters away from infected cells can reach an antiviral state before a virus arrives. In summary, IFN response by cells at low MOI differs from that of cells in high MOI infection for two reasons: 1) at high MOI, infection starts synchronously in most cells, and paracrine IFN signaling is less relevant since neighbors are already infected; 2) at low MOI, infection is spatially sparse and paracrine signaling precedes viral arrival, so cells have time to become virus resistant. Because the typical travel time of viruses is of similar order of magnitude to the typical 48h of viral load peak and because the typical distance between infected cells influences the total ISG expression, we argue that, for low MOI IAV infection of the *in vitro* epithelium, spatial structure is an important aspect of infection dynamics. In this simple argument, we do not consider transport by advection, for example, caused by cilia movement. Other studies highlight the importance of spatial structure for IAV spread (41–44), emphasizing the locality of cell signaling and viral transmission, as well as heterogeneity in cell responses and cell density.

In addition to incorporating spatial structure, we took an **Agent-Based Modeling (ABM)** approach, which considers each cell as a system with its own parameters and states. This is an appropriate strategy to capture tissue effects that emerge from individual cell behavior and heterogeneity of cell behavior. When coupled to spatial structure, the agent-based approach can represent cell-to-cell interaction in detail (e.g., paracrine IFN signaling, effects of different types of SIPs in cells, and viral genome complementation).

Finally, our model explicitly considers the effects of antiviral responses on bystander cells (i.e., immediate neighbors to the infected cells). A bystander cell exposed to IFN that acquires an antiviral state before infection by a virion has a different fate from a cell infected by a virion before IFN exposure. The effectiveness of NS1 and therefore the cell antiviral state, as well as the rates of spread in the environment for virus and IFN, which we will treat as diffusive with constant diffusion coefficients, determine the outcomes of the infection. The outcomes of complementation of viral genes in infected cells depend on which genes are nonfunctional in each infecting SIP and how long the cell was infected before complementation. Since cell fates depend on the combination of productive viral genes from all the infecting virions, a predictive model of infection and anti-viral resistance must explicitly track nonfunctional viral genes in each individual cell, a goal which is compatible with the agent-based approach but cannot be readily achieved with ODEs or PDEs. Although the rates of viral release, IFN release, and the concentrations of proteins vary between individual cells, our experimental data only measure population-averaged IFN and viral levels. We simplify this variability by modeling a population of cells in which IFN and virus are released at constant rates, and in which proteins are treated as either present or absent in each cell.

To our knowledge, this is the first study incorporating a spatial ABM of IAV infection with both the molecular mechanisms underpinning viral antagonism and viral complementation of SIPs. To reproduce observed IFN levels across different MOIs, we developed a spatial ABM of the *in vitro* cell culture that integrates **Boolean stochastic network** (**BSN**) models of viral-host antagonism, viral replication, and IFN response. It also includes spatial diffusion of virions and IFN, SIPs, and viral complementation. This framework allows us to assess the MOIs for which simplified models that omit SIPs can adequately predict time series of IFN levels.

Response to a given strain of IAV infection *in vivo* varies between individuals (45), however, *in vitro* experiments often employ **c**ells generated from a single donor. If the cell-level variability between individuals is substantial, experiments using a single cell line and simulations developed and calibrated from them might not predict outcomes for individuals with different genetic profiles and personal histories. In scenarios where different groups of people present significant differences in immune response, e.g., age groups, sex, and immunosuppressed patients, the correct approach is to calibrate the model once per group instead of the population average. We therefore quantified the differences in temporal responses to infection in cultures derived from a representative variety of donors. However, as we will discuss below, our experiments did not reveal variation significant enough to require recalibration of our computational models to the individual donor cell lines.

## Materials and Methods

### Cell Culture

Madin-Darby canine kidney overexpressing sialic acid (**MDCK-Siat**) cells were cultured in Dulbecco’s Modified Eagle Medium (**DMEM**, Sigma-Aldrich) with 10% fetal bovine serum (**FBS**, Gibco Life Technologies), and 100 U penicillin/mL with 100 μg streptomycin/mL (Quality Biological) at 37°C with air supplemented with 5% CO_2_. Infectious medium for IAV (**IM**) was used in all infections and consisted of DMEM with 5 μg/mL N-acetyl trypsin (**NAT**), 100 U/mL penicillin with 100 μg/mL streptomycin, and 0.5% bovine serum albumin (**BSA**) (Sigma).

Human nasal epithelial cells (**hNEC**) (Promocell, lots 489Z024.3, 502Z025.1, 499Z012.1, 493Z033.1, 502Z024.2) were grown to confluence in 24-well Falcon filter inserts (0.4-μM pore; 0.33 cm^2^; Becton Dickinson) using PneumaCult™-Ex Plus Medium (Stemcell). Donor information is listed below:

| Lot | Age | Sex | Ethnicity | PDT [h/PD] | Viability [%] | Smoker Status | Asthma J45 | COPD J44 Group | COPD J44 Stage |
| --- | --- | --- | --- | --- | --- | --- | --- | --- | --- |
| 489Z024.3 | 57 | FEMALE | CAUCASIAN | 32.5 | 74 | NON-SMOKER | NO | GROUP B | STAGE II |
| 502Z025.1 | 60 | MALE | CAUCASIAN | 29.3 | 87 | NON-SMOKER | NO INFORMATION | NO | NO |
| 499Z012.1 | 76 | MALE | CAUCASIAN | 26.3 | 84 | NON-SMOKER | NO | NO INFORMATION | STAGE II |
| 493Z033.1 | 55 | FEMALE | CAUCASIAN | 51.8 | 86 | NON-SMOKER | NO INFORMATION | NO | NO |
| 502Z024.2 | 51 | FEMALE | CAUCASIAN | 35.6 | 88 | NEVER-SMOKER | NO | NO INFORMATION | NO |

Confluence was determined by a transepithelial electrical resistance (**TEER**) reading above 250Ω by Ohm’s law method (8,46) and by examination using light microscopy and a 10X objective. The cells were then differentiated at an air-liquid interface (**ALI**) before infection, using ALI medium as basolateral medium as previously described (47,48). Briefly, both apical and basolateral media were removed and ALI differentiation media (Stem Cell Technologies, Pneumacult ALI Basal Medium) supplemented with 1X ALI Maintenance Supplement (StemCell Technologies), 0.48 μg/mL hydrocortisone solution (StemCell Technologies), and 4 μg/mL Heparin sodium salt in phosphate buffered saline (**PBS**) (StemCell Technologies) was replaced on the basolateral side only. Fresh media were given every 48 hours. We will refer to the differentiation media as ALI media. Once mucus was visible, apical washes were performed weekly with PBS to remove excess mucus. Cells were considered fully differentiated after 3 weeks and when cilia were visible using light microscopy using a 10X objective. All cells were maintained at 37°C in a humidified incubator supplemented with 5% CO_2_.

### Virus Seed Stock and Working Stock Generation

Influenza A Virus (A/Baltimore/JH-22377/2022 (H1N1), GISAID EPI_ISL_17626298), was isolated from samples obtained through the Johns Hopkins Hospital network as part of the CEIRS network (49). To generate virus working stocks, MDCK cells were seeded in a T150 flask and infected at a MOI of 0.001 with virus diluted in IM. After one hour, the inoculum was removed, and fresh IM was added. When a cytopathic effect was seen in approximately 50% of cells, the supernatant was harvested, aliquoted, and stored at −65°C.

### TCID50 Assay

MDCK-Siat cells were grown to 70–90% confluence in 96-well plates. After being washed twice with PBS+, ten-fold serial dilutions of the viruses in IM were made and each dilution was added to 6 wells. The plates were incubated at 37°C with 5% CO_2_ for 4 days. The cells were fixed by adding 75 μL of 4% formaldehyde in PBS per well for at least 4 h and then stained with Naphthol Blue Black solution for 1 hour. Endpoint values were calculated by the Reed-Muench method (19).

### Viral Infections

For ALI infections, an MOI of 0.1 per cell was used. The apical side of the transwell was washed 3 times with IM, with a 10-minute incubation at 37°C between each wash. The virus inoculum was diluted in IM and 100 μL was added to the apical side of cells and allowed to incubate for 2 hours. The inoculum was then removed, the cells washed 3 times with PBS and returned to the incubator. At the indicated time points, a 10-minute apical wash was performed on wells with IM and basolateral media was collected and stored at −80°C. Infectious virus particle production in apical washes was quantified using TCID50 on MDCK-Siat cells.

### Immunoprofiling

Interferon panels (PPX-MXRWHCG, lot 449667-000, Luminex Corporation) were run as recommended by the manufacturer. Briefly, samples were thawed and centrifuged at 10,000 g for 10 minutes to pellet any particulates. Standards were reconstituted in 50 μL of ALI media, briefly vortexed, and incubated on ice for 10 minutes. The plate was prepared by adding 50 μL of Capture Bead Mix to each well and washing with 150 μL 1x PBS wash buffer. A 50 μL aliquot of each standard was added to the plate in duplicate followed by single technical replicates of each experimental sample (3-4 biological replicates). The plate was then sealed and placed on a shaker overnight at 600 rpm at 4°C. The following morning, the plate was shaken for an additional 30 minutes at room temperature and then washed twice. 25 μL of Biotinylated Detection Antibody Mix was added to each well and the plate was once again sealed and shaken at 600 rpm for 30 minutes at room temperature. The plate was washed twice and 50 μL of Streptavidin-PE was added to each well, and the plate was shaken at 600 rpm for 30 minutes at room temperature. The plate was washed twice and 120 μL of reading buffer was added to each well. The plate was shaken one last time at 600 rpm for 5 minutes and then was read on the Luminex machine.

### IFN Decoupling

IFN-lambda (PBL Biosciences) working stocks were prepared in PBS with 0.1% BSA and stored at −80°C. Fresh aliquots were taken for each treatment. For IFN decoupling experiments, cultures were pretreated with 500 U/mL of IFN-λ added to the basolateral ALI media 24 hours prior to infection. Basolateral media was collected every 24 hours and stored at −80°C. Immediately following each collection, the media were replaced with fresh media containing IFN-lambda. Infections were performed as described above.

### Simulation Methods

Our conceptual model of IAV infection and IFN response in individual cells seeks to account for two key biological observations. **1)** The NS1 viral protein, a potent IFN signaling suppressor within individual cells, efficiently inhibits IFN production and IFN signaling across the cell population in the culture at high MOI; however, **2)** immune-competent cell cultures can still exhibit robust population-level IFN response at low MOI. Our model is structured with the following assumptions:

1. Cell behaviors depend on whether they have been exposed to IFN and whether they have been infected by IAV:

a. Non-infected cells in a low IFN concentration environment (**N**on-Infected **L**ow-IFN **C**ells - **NLCs**): produce and secrete IFN at a low baseline rate and are susceptible to infection.
b. Non-infected cells in a high IFN concentration environment (**N**on-Infected **H**igh-IFN **C**ells - **NHCs**): gain protection from infection by activating ISGs, and are less susceptible to infection than NLCs.
c. Infected cells in a low IFN concentration environment (**I**nfected **L**ow-IFN **C**ells - **ILCs**): may produce and secrete IFN if IFN mRNA is produced before NS1. If all viral genes are present and functional, ILCs eventually start releasing virions and keep doing so until they die.
d. Infected cells in a high IFN concentration environment (**I**nfected **H**igh-IFN **C**ells - **IHCs**): tend to produce and secrete larger amounts of IFN than ILCs when IRF7 is active in response to external IFN. If all viral genes are present and functional, IHCs eventually start releasing virions and keep doing so until they die.
2. Cells acquire antiviral resistance behaviors by sensing IFN concentration in the medium and activating transcription of ISGs. The higher the extracellular concentration of IFN, the faster the activation of ISGs. We consider four classes of proteins produced by ISGs: 2′-5′-oligoadenylate synthetase (OASs) and protein kinase R (PKR) inhibit production of viral proteins, interferon-induced transmembrane proteins (IFITMs) reduce virus endocytosis, and interferon-regulatory factor 7 (IRF7) boosts IFN production when a resistant cell is infected. ISG transcription decays after initial activation.
3. Most virions have at least one damaged gene incapable of coding for its corresponding viral protein, i.e., most virions are SIPs (28).
4. Each viral gene has an associated defect probability, and defect probabilities are independent. We estimate the defect probability of a gene assuming a constant probability of loss-of-function point mutation per base pair. The probability *P* =(1 - *P_db_*)*^S^*that an individual gene is defective depends on its size *S*, and on the probability *P_db_* that a single base defect is fatal for the protein function. More details in the Supplemental Materials Section 1.
5. Individual cells can endocytose multiple virions during the early phases (first 5 hours) of infection (i.e., “superinfection”). Cells releasing viruses are incapable of further superinfection.
6. A cell infected only by SIPs with defective NS1 gene(s) will secrete IFN.
7. A cell containing functional NS1 protein does not transcribe new IFN mRNA. NS1 also blocks activation of ISGs.
8. A cell that synthesizes IFN mRNA before NS1 is produced will initially secrete IFN. Since IFN mRNA decays with a half-life of about 12 hours (50–52), NS1 will eventually halt IFN synthesis. The time scale for NS1 production is 4h after infection, while the time scale for IFN mRNA production is 4h after infection.
9. A cell infected by an SIP lacking one or more of the genes necessary for viral genome replication (PA, PB1, PB2, NA) will not produce virions.
10. A cell infected by an SIP lacking one or more of the genes necessary for viral transport, binding, and release (NA, HA, NEP, M1, M2) will not release virions.
11. SIPs can complement their genomes inside a cell upon superinfection, potentially rescuing NS1 antagonism, viral genome replication, and virion release.

We embody these mechanistic assumptions in a computational model to simulate the time course of IFN levels in an *in vitro* epithelial cell culture after infection by IAV. Supplemental Table 1Error: Reference source not found lists all model assumptions with their motivations and references.

For simplicity, this model does not account for several biological components and mechanisms, notably:

1. We do not consider DIPs.
2. Gene defects arise from fatal point mutations in viral RNA. Biologically, SIPs originate from a variety of defect sources, such as the loss of one or more full segments (53). In our model, we consider only fatal point mutations as sources of gene defects.
3. Gene defects are independent within a virion. Biologically, a defective gene can compromise transcription of other genes in the same segment.
4. All virions have the same gene defect probabilities, regardless of which cell produced them. Biologically, cells infected by non-infectious particles are more likely to release non-infectious particles than cells infected only by IUs.
5. We represent viruses as a continuous density field rather than as individual agents.
6. We do not consider minimal time delays due to virion internalization and unpacking.
7. We do not consider minimal time delays during protein and RNA synthesis arising from transcription, translation, and transport.
8. The virtual cell culture is 2D and includes a single type of epithelial cell. All cells have identical parameters.
9. We do not explicitly model the removal of viruses from the culture due to culture wash events for quantification.
10. Virus endocytosis and internal cell states are Boolean and transitions between different state levels are stochastic with constant probability rates that are the same for all cells, but different for different processes.
11. Levels of viral proteins in the cell are Boolean (proteins are either present or absent).
12. We do not consider IFN-induced apoptosis and NS1-induced suppression of apoptosis.
13. We do not consider epithelial cell proliferation and tissue recovery. While we have explored these and observed oscillations in the viral load time series (data not shown), they are not the focus of this work.

See Supplemental Table 2 for more details about these simplifications.

We implement the mathematical model as a computer simulation using the open-source CompuCell3D (CC3D) (20) Virtual-Tissue modeling framework. The *in vitro* environment is represented by a 2D square lattice, containing the cells and fields. Each cell contains a BSN expressed in the MaBoSS language (21). The nodes in the network represent proteins and the activity of key processes like virus production. The state of the network evolves according to continuous-time stochastic Boolean dynamics, with transition rates governing node activation and deactivation.

The simulation begins with the cells in a regular square grid. We then run an 8-hour relaxation during which cell borders fluctuate following Cellular Potts Model/GGH-dynamics (20), allowing cells to relax to a hexagonal configuration. Because cells have a baseline IFN secretion rate during relaxation, nodes of the BSN related to ISGs also adjust during this period. After relaxation, we reset the simulated time to zero, add virus to the extracellular environment, and initiate infection.

We represent the extracellular virions as a continuous density field, where one unit of this field in one lattice site is equivalent to one virion. After relaxation, we seed viruses by adding one unit of virus field in a random lattice pixel (See Supplemental Table 3 for more details on virus seeding). The MOI parameter and the total number of cells determine the initial total virus count supplied.

The computational model contains viral, intracellular, and culture-level parameters, such as secretion rates, diffusion coefficients, upregulation and downregulation probability rates, and gene defect probabilities, as well as parameters that control the initial conditions, such as MOI, lattice size, and total number of cells. Some parameters we estimate from literature; others we assign physiologically reasonable values. See Supplemental Table 3 for parameters and estimates. To fit the experimental data, we allow three parameters to vary: MOI, IFN secretion rate, and virus release rate. Calibration of these three parameters is sufficient to match available experimental time series without adjustment of the remaining parameters. The simulation is very sensitive to these free parameters, with virus release rate and IFN secretion rate influencing viral load and IFN levels, and MOI influencing the time scale of the infection. We did not find published estimates of virus release rate and IFN secretion rate per infected cell in *in vitro* assays. Although our calibrated simulation reproduces experimental time series for the IFN level and viral load, it failed to reproduce experimental NS1 expression in infected cells over time (2).

We calibrate the parameters for the per-cell virus release rate, IFN secretion rate, and MOI with experimental time series to constrain our parameter values and acquire a Rough Order of Magnitude (**ROM**) estimate of the parameters. We utilize time series of IFN level, in units of [µg/mL], and viral load, in units of virus count per cell because they reveal both magnitude and time scale of extracellular virus and IFN. By trial and error, we identified a candidate parameter set that provided reasonable agreement with the experimental time series. We then did a Monte Carlo parameter exploration around these candidate parameter values to constrain the region of parameter space near our initial guess and plotted error threshold regions (Supplemental Figure 2), defined as the sets of parameter values around the candidate parameter set that yielded an error below a specified cutoff. The error in a single time series is the sum of the absolute values of the residuals. To fit the viral load and the IFN time series simultaneously, we calculate the total error as the sum of the normalized errors for each time series. We normalized each error contribution by the corresponding maximum experimental value, providing a simple, scale-invariant measure that allows direct comparison across variables with different magnitudes. We then sampled parameter combinations around the candidate set and visualized the resulting regions. These plots illustrate the degree of practical identifiability of the parameters and highlight correlations among them. To test robustness, we constructed allowed parameter regions for two error thresholds. Parameter regions should shrink in all parameter directions as the tolerance decreases and the allowed region should stay simply connected. We found that the stricter threshold resulted in a single, very small, allowed region of parameter space, and, surprisingly, our initial guess was at the center of the allowed region for the smaller threshold. Thus, the three parameters are highly identifiable, and our initial guess was very close to the global optimum. We therefore accepted our guess as our reference parameter set for our remaining analysis.

To understand how each remaining parameter affects the simulation, we ran a single factor Local Sensitivity Analysis (**LSA**). We increased each parameter individually by 50% while the other parameters remained at their default values, see Figure 3. Parameters were also decreased by 50% (data not shown). We measured the area under the curve of IFN concentration and viral titer relative to default parameters for high/low MOI. We averaged over 200 simulation replicates for each parameter set.

We needed tens of thousands of replicates to generate data for the LSA and the bootstrap analysis. Therefore, we ran simulations *en masse* on a small Linux HPC cluster consisting of four nodes, each with 24 subnodes and 96 GB of memory per node. Each low MOI simulation takes about 5 minutes to run on this cluster; high MOI simulations run faster.

Source code, parameters, simulation results and simulation movies are available on Zenodo [https://doi.org/10.5281/zenodo.20303705] and on GitHub [https://github.com/pdalcastel/IAV_Infection_CC3D_MaBoSS_2026]. See the Supplemental Information for additional details.

## Results

### Donor-to-donor heterogeneity plays only a minor role in IFN levels in the culture

To characterize the effect of donor heterogeneity and calibrate our computational model, we measured viral load and IFN time series for *in vitro* cultures using cells sourced from different donors, in response to viral infection and IFN treatment. Cultures from all donors showed broadly similar responses to H1N1 infection, with comparable viral replication kinetics and peak titers (Figure, Panel A). Statistically significant differences included higher virus production in Donor 5 than in Donor 1 at 12 hpi, and in Donor 2 compared with Donor 3 at 48 hpi. However, these differences were not sustained over the course of the experiment.

Cultures from all donors also showed similar time series for expression of 11 pro-inflammatory cytokines, with overall cytokine production increasing over time (Figure, Panel B). We also observed initial inhibition of IL-29 (also known as IFN-λ), consistent with the activity of NS1. The greatest differences observed between donors occurred at 12 hpi; however, the cytokines are also produced at low rates at this time, meaning small changes can have large effects on these values. These data support calibrating the model to the average of viral load and IFN levels across all donors, rather than treating each donor as a single model calibration case.

To look for donor-based variability in sensitivity to IFN-mediated inhibition of viral replication, we pre-treated cultures from the five donors with IFN-λ starting 24 hours prior to infection and compared the resulting virus production time series to those from cultures that received vehicle only (Figure, Panel A, green triangle data points). As expected, IFN-λ pre-treatment significantly inhibited viral replication in all samples. Viral replication kinetics or peak titer were statistically indistinguishable across cultures from all donors undergoing the same treatment.

We used the *in vitro* viral load and IFN concentration time series, averaged across donors, to calibrate the model parameters, which included IFN secretion rate, virus release rate, and MOI. We estimated an IFN secretion rate per cell per hour of 4.55 [µg•mL^-1^cell^-1^h^-1^] (considering 0.1 mL of sample volume and 10^7^ cells). This rate estimate may be useful to parameterize future mechanistic models that implement IFN secretion per cell and attempt quantitative validation of single-cell IFN response. We estimated a virus release rate of 1.9 *×* 10^3^ virions per cell per hour (this rate includes SIPs as well as IUs). Finally, we estimate an MOI of 1.2 *×* 10^-3^, *i.e*., around 1 IU (fully infectious particle) every 1000 cells. Monte Carlo exploration of the parameter space reveals that, under a 10% error threshold, acceptable fits were obtained for IFN secretion rate in the [4.4, 5] range, virus release rate in the [1.6, 2] *×* 10^3^ range, and MOI in the [0.8, 1.6]*×* 10^-3^ range (Supplemental Figure 2).

The slope of the simulated viral load curve in the period between 24 hpi and 48 hpi does not perfectly match the *in vitro* experiment. The simulation predicted that viral load increase starts sooner than it does experimentally and saturates later. This deviation is possibly due to omission of transport and transcriptional/translational delays in the BSN, leading to virus release and IFN production earlier than expected. Increasing average transition times improves fit at earlier times but leads to mismatches at later times. The best fit of a BSN cannot perfectly match both the onset and the end of the viral load curve. For the same reason, our model fails to predict both onset and saturation times in the time series of total NS1 expression, particularly Figure 8 Panel B of reference (2) by Ramos *et al*. These data show null NS1 expression until 3 hpi, indicating that time delays are present between virus internalization and NS1 production. Since the state of the NS1 node in the BSN transitions stochastically according to a constant probability rate, we can only capture the average activation time, not the start of activation nor the finish. Finally, the model predicts that IFN treatment prior to infection decreases viral load area under the curve, with higher levels of IFN leading to larger decreases. Supplemental Movie 2 shows the effect of IFN treatment on the plaque dynamics and IFN levels.

### Intermediate MOIs show the highest fraction of IFN mRNA expressing cells, NS1 deletion recovers high IFN levels at high MOI, and running the simulation without SIPs fails to reproduce experimental results

After calibrating the simulation to our experimental data, we compared simulation predictions of IFN levels, the fraction of IFN expressing cells, and the number of viruses released per cell to literature data not used in the construction of our computational model.

The computer simulation showed strong NS1-induced inhibition of IFN production at high MOI, high IFN production at low MOI, and some IFN production in the presence of NS1-incompetent virions, even at high MOIs (Figure, Panel A). This result aligns with work by Marcus P. (25) and work by Tawaratsumida K. and colleagues (26), where they compare IFN levels induced by NS1-incompetent vs. wild-type viruses. Work by Yang Q. and colleagues and Russel A. also indirectly supports the observation that NS1 is a potent inhibitor of IFN production and signaling (20,23).

IFN levels at 24 hpi are highest for an intermediate MOI, with both lower and higher MOI leading to lower IFN levels (Figure, Panel C). This trend is also observed when measuring the number of cells expressing IRF7 at 24 hpi. Alalem M. and colleagues reported a similar trend in the relative gene expression of MxA in figure 1c (24). MxA is an antiviral protein derived from an ISG. Although we do not explicitly model MxA, we use IRF7 as a proxy, as IRF7 is also derived from an ISG. This trend of peak IFN response at intermediate MOIs disappears when we set all gene defect probabilities to zero, i.e., all virions are IUs, and there are no SIPs. For the comparison to be fair, we calibrated the simulation without SIPs to the same experimental IFN and viral load time series as the original simulation with SIPs, so the parameter values of virus release rate and IFN secretion rate are different from the simulation with both IUs and SIPs.

**Figure 1.**
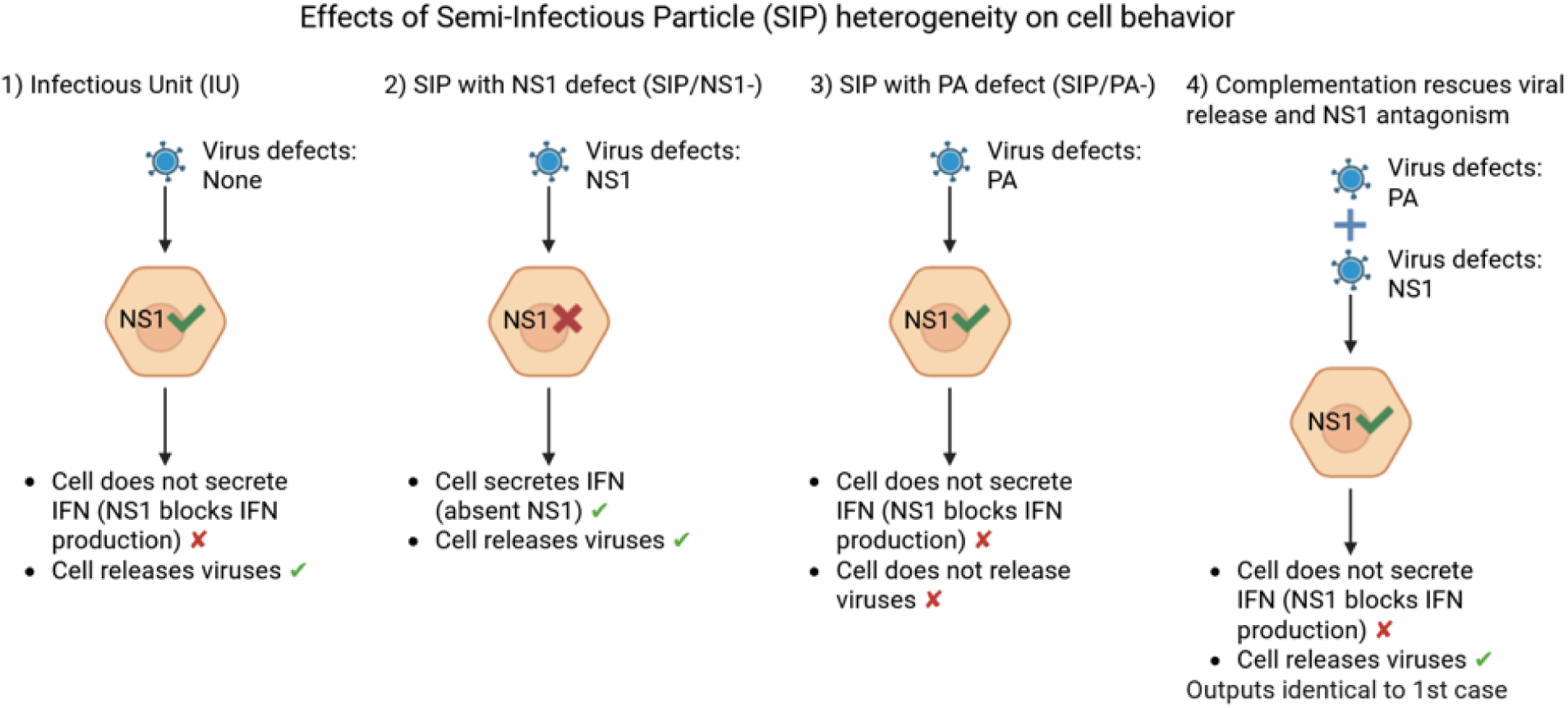
Simplified conceptual model of genome complementation during coinfection by SIPs and the outcomes for different patterns of missing functional proteins. Biologically, the outcomes depend on the type of gene defect, while our simplified conceptual model assumes that defects in viral genes result from independent fatal point mutations. Here we consider defects of NS1 and PA (one of the subunits of the viral RNA polymerase), which belong to different segments of the viral genome. A functional NS1 gene inhibits interferon production, while a functional PA gene is required for viral replication, and therefore required for releasing virions. 1) A cell infected by an IU with no defects expresses all viral proteins, preventing IFN production and allowing release of infectious viral particles. 2) A cell infected by an SIP with a defective NS1 gene produces IFN and releases infectious viral particles. 3) A cell infected by a SIP with a defective PA gene and a functional NS1 gene does not produce IFN and does not release infectious viral particles. 4) A cell multiply infected by two SIPs, one with a defective PA gene and the other with a defective NS1 gene behaves like an infection by an IU, inhibiting IFN production and allowing release of infectious viral particles.

NS1 antagonism and complementation are key to interpreting the previous result. At high MOI, complementation is frequent, increasing the presence of productive NS1 in infected cells. At an intermediate MOI, cells take longer to achieve full viral genome complementation, and if NS1 is absent, these cells will produce IFN during this time. The lower the MOI, the longer the infection takes to spread; therefore, intermediate MOI is optimal to observe peak IFN response at 24 hpi. Previous computational models of IAV infection have examined the complex viral life cycle and virus–host interactions (36,37,44,54–62), and the physiological effects of DIPs (63–66) and SIPs (29). However, to our knowledge, no model addresses the relationship between NS1 antagonism, gene defects in SIPs and the rescue of NS1 antagonism via complementation. Instead, models aggregate parameters (such as reducing effectiveness of NS1 antagonism) to capture the overall effect of SIPs. These models can reproduce time series of IFN for specific MOI regimes but do not capture the dependence of IFN levels on MOI.

Virus release rates vary between infected cells, with a minority of cells responsible for most released viruses (Figure, Panels D and E). The linear-scale (**D**) and log-scale (**E**) distributions of the total number of viruses released per cell are similar to the distributions reported by Heldt F. et al. in their Figure 1b (67), and Kupke S. et al. in their Figure 2a, (68), respectively. The same pattern occurs with IFN, where a minority of cells produce and release most IFN (data not shown). With our simulation, we can measure the time elapsed from first viral entry to full genome complementation per cell, showing that cells frequently take longer than 5 h to achieve full genome complementation (Figure, Panel F). During these 5 h, cells that specifically lack a functional NS1 gene have enough time to produce and secrete IFN. In a study by Brooke et al., Fig. 6 shows a case of low MOI (<0.01) IAV infection at 15 hpi with two plaques of roughly 10 – 20 cells in diameter, and around 20 infected cells that failed to start a plaque due to lack of a complete viral genome (23). It would take multiple extra rounds of virus replication for these cells to receive new infectious virions and achieve full complementation, i.e., longer than 15 h. Although our histogram captures this observation, we did not find an equivalent experimental histogram in the literature. Obtaining a similar histogram from experiments would be challenging, highlighting an advantage of this simulation. We also provide a time series of the fractions of fully and partially infected cells (Figure, Panel F). These series show that most infected cells lack at least one viral gene in the earlier phases of infection until 20 hpi (low MOI scenario). In the later stages of infection, most cells become fully infected.

**Figure 2.**
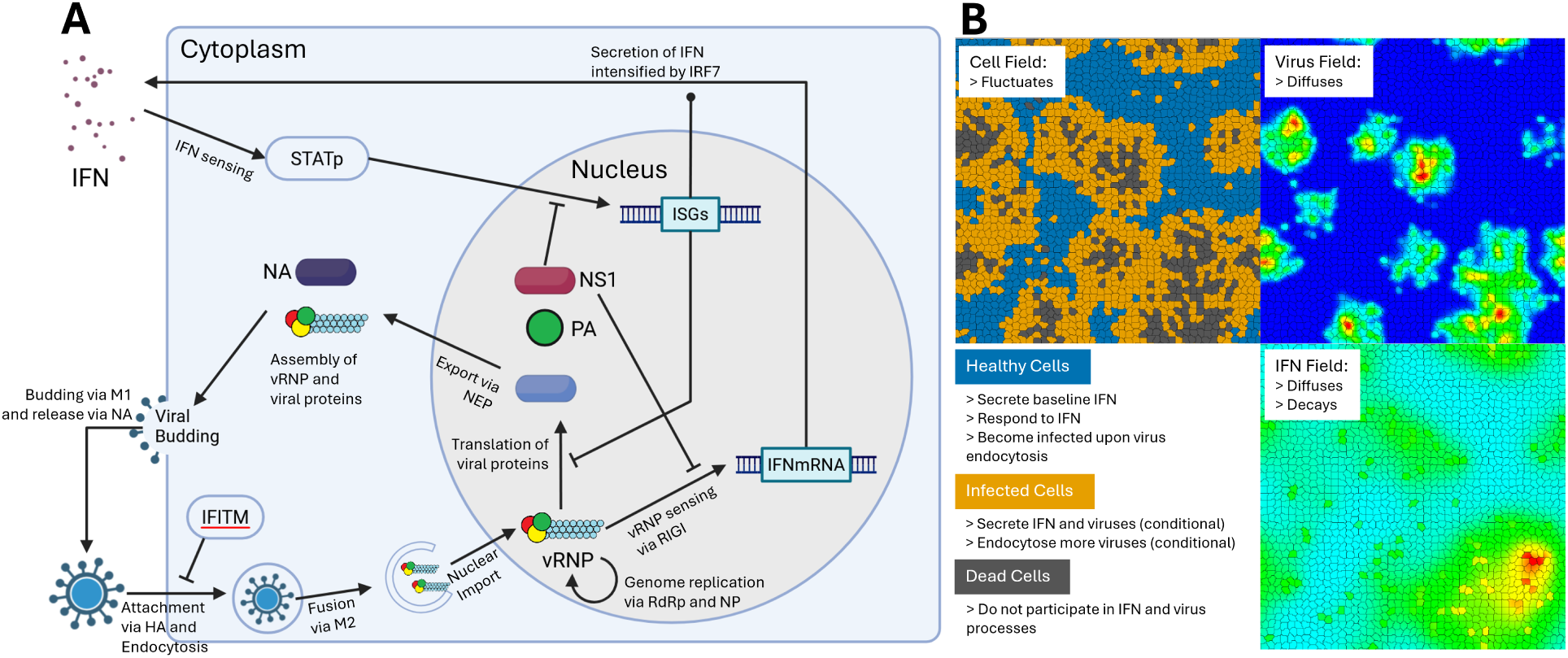
Subcellular signaling and response network. **A)** Conceptual model of the intracellular pathways governing viral-host-cell interactions, featuring viral internalization, virus sensing by the cell, viral replication, IFN production, IFN sensing, activation of ISGs, suppression of viral functions by ISG proteins, and suppression of cell immunity by the viral protein NS1. In our computational model, each cell agent contains a BSN, which includes key nodes and interactions that represent the main components of the conceptual model, implemented using the MaBoSS framework. See Supplemental Figure 1 for a code-level diagram with all nodes and logic operators. **B)** The agent-based tissue includes healthy, infected and dead epithelial cell objects on a 2D lattice, implemented in the CompuCell3D framework. Simulated cells interact with the extracellular concentration fields of virus and IFN. Cells’ behaviors depend on their state (healthy, infected, dead) and internal network state.

**Figure 3.**
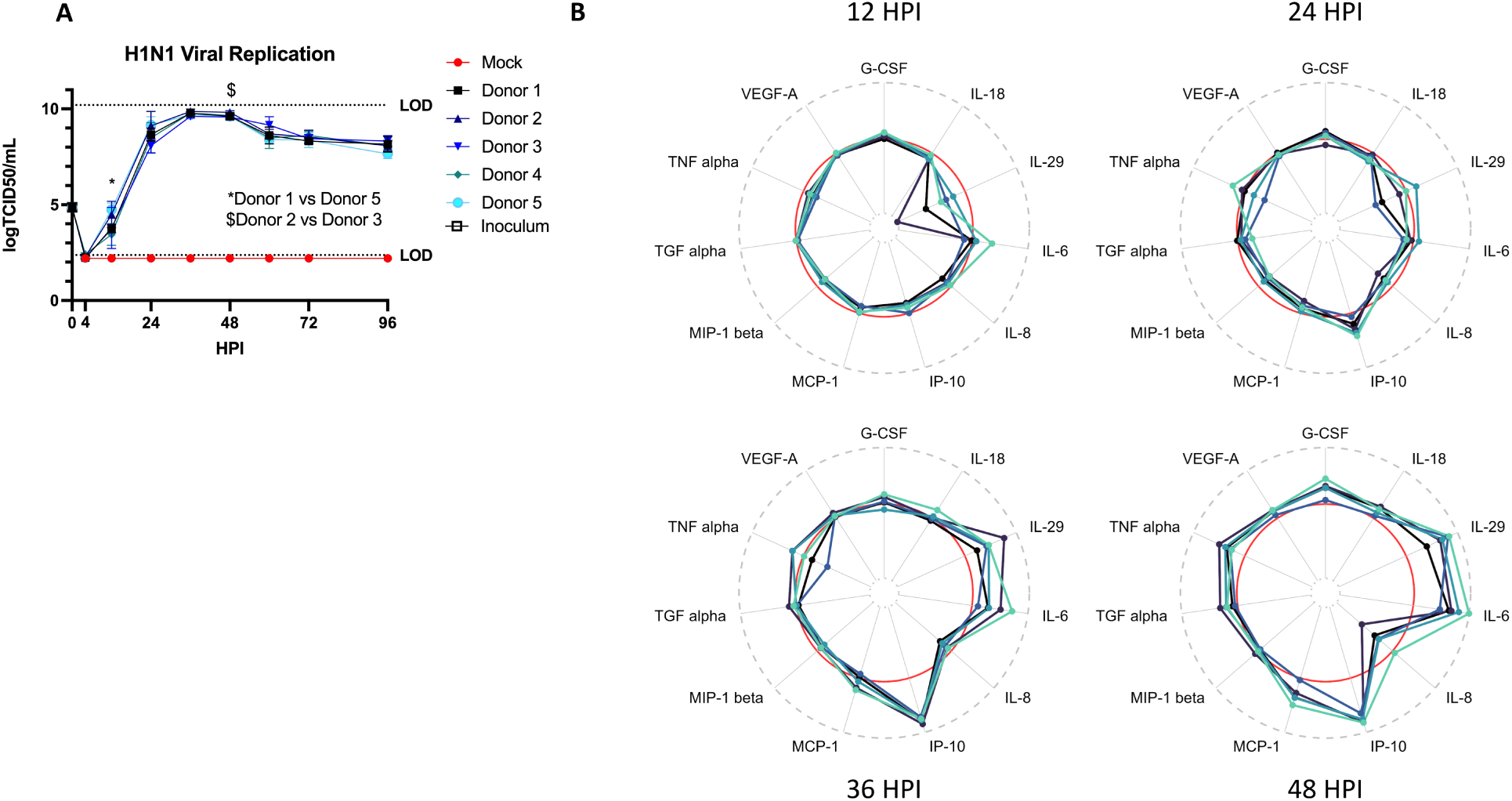
Comparison of IAV H1N1 replication and epithelial cell innate immune response in hBECs derived from different donors. A) Low MOI multi-step H1N1 growth curve; n= 4 wells per donor per condition (Mock or infected). *p < 0.05 (two-way repeated measures ANOVA with Tukey’s posttest, only 12–48 h included). Dotted line indicates limit of detection. B) Basolateral secretions of various immune molecules at the indicated timepoints after low MOI inoculation. Log fold-change relative to matched mock-infected wells collected at the same time is shown for each donor set. The red line indicates log fold change of 0 (or no difference). Values outside the red circle indicate higher expression in H1N1-infected wells, while values inside the red circle indicate higher expression in mock-infected wells. *p < 0.05 (two-way repeated measures ANOVA with Tukey’s posttest on measured values).

**Figure 4.**
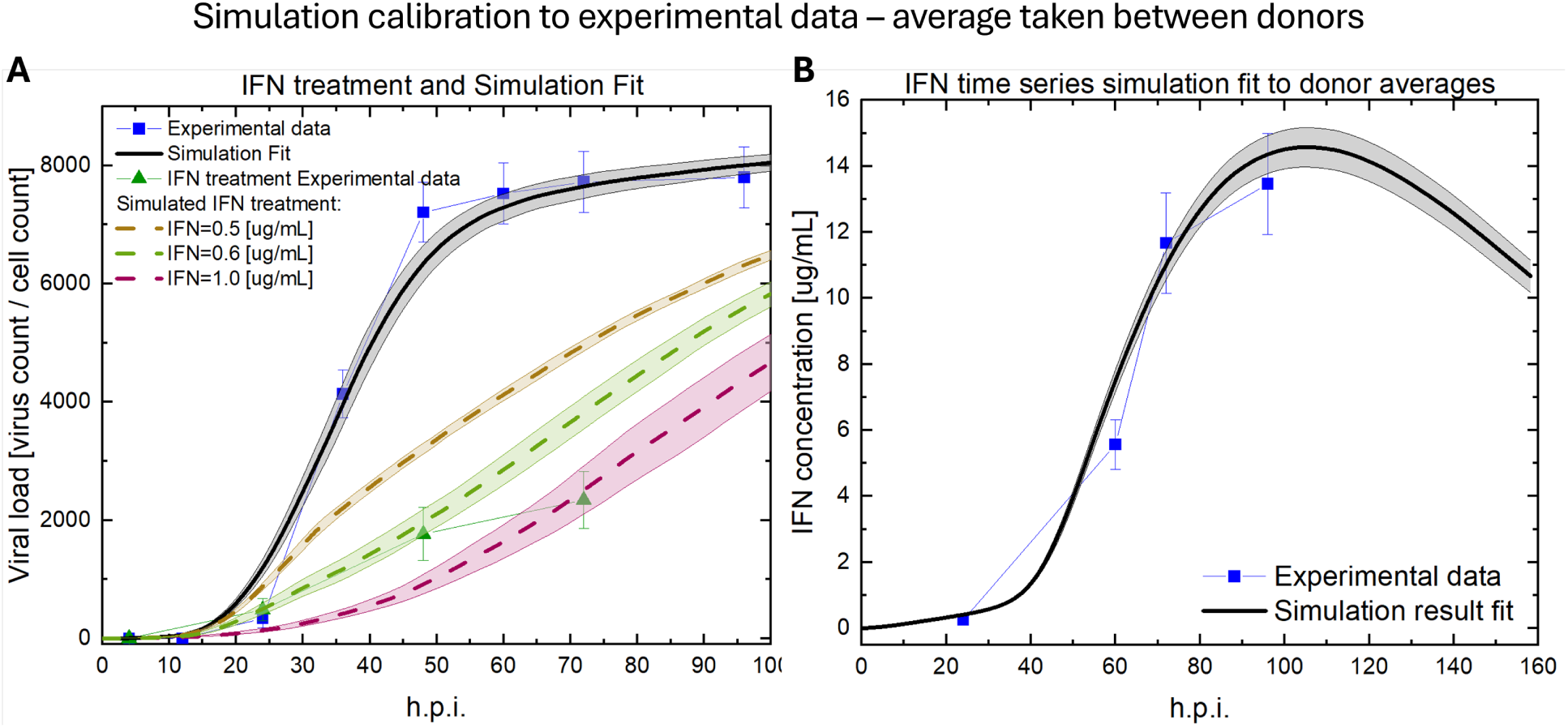
Calibration of the simulation to experimental time series for viral load and IFN concentration with IFN secretion rate, Virus release rate, and MOI used as fitting parameters. Connected points represent experimental data, solid lines represent the simulation result with the candidate reference parameters, and dashed lines show simulation results after IFN treatment. The grey error band illustrates the standard error of the mean derived from 10 independent simulations. **A**) Viral load from experiment and simulation. **B**) IFN level from experiment and simulation.

**Figure 5.**
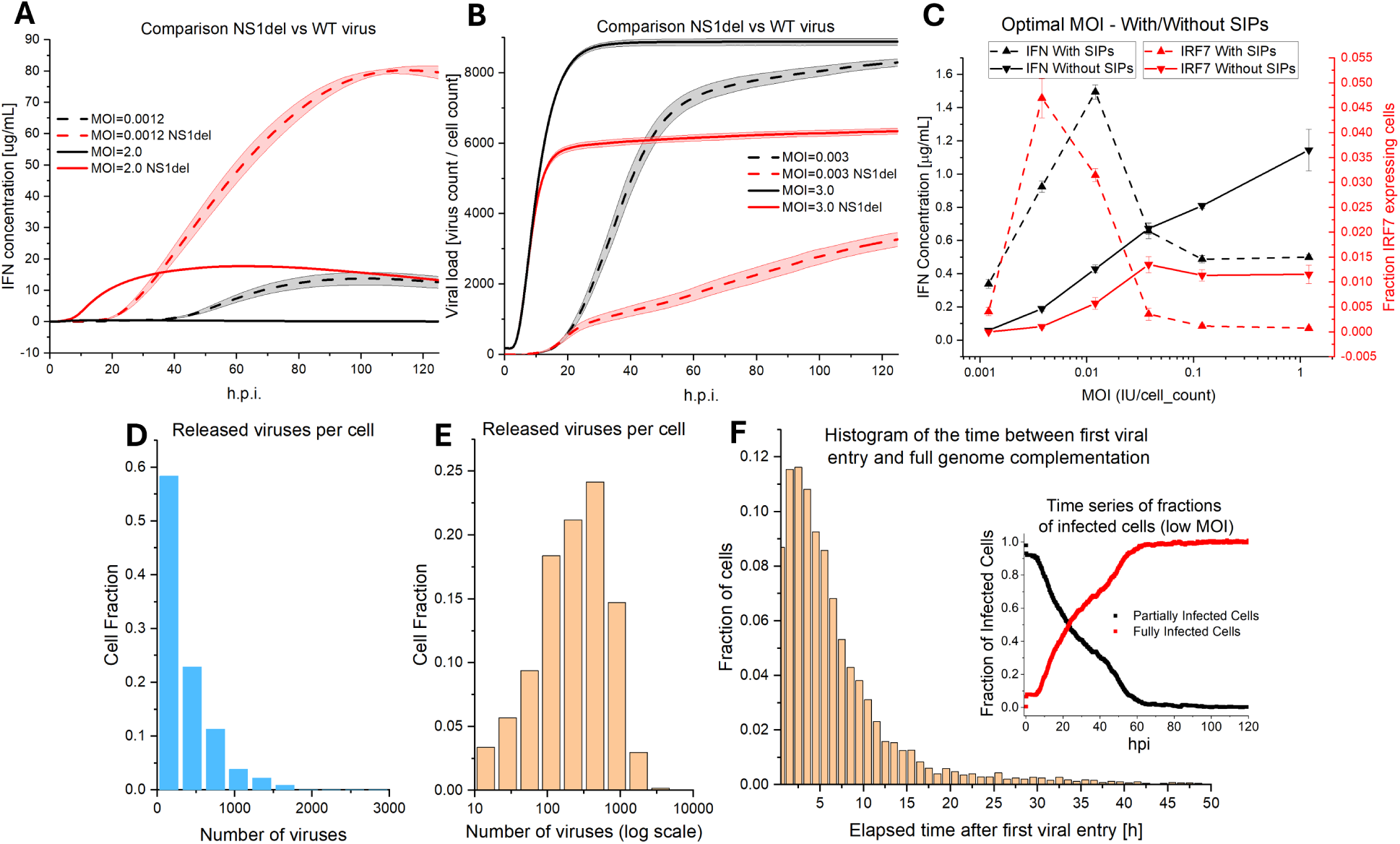
The calibrated computer simulation shows substantial IFN production at low MOI and IFN suppression at high MOI. **A)** Time series of IFN levels for NS1del (NS1-incompetent virus), WT (wild type), and high/low MOI. IFN levels increase for NS1del virus and lower MOI. **B)** Time series of viral load for NS1del (NS1-incompetent virus), WT (wild type), and high/low MOI. **C)** Levels of IFN and fraction of cells expressing IRF7 at 24h peak at an intermediate MOI, in agreement with experimental results. This peak vanishes when we remove SIPs. **D, E)** Histograms of the fractions of cells that released a certain number of viruses in linear (**D**) and logarithmic (**E**) scales. **F)** Histogram of the time elapsed between first viral entry and full viral genome complementation in a single cell throughout a low MOI simulation. The embedded graph shows time series of the fractions of infected cells for the same MOI.

**Figure 3.**
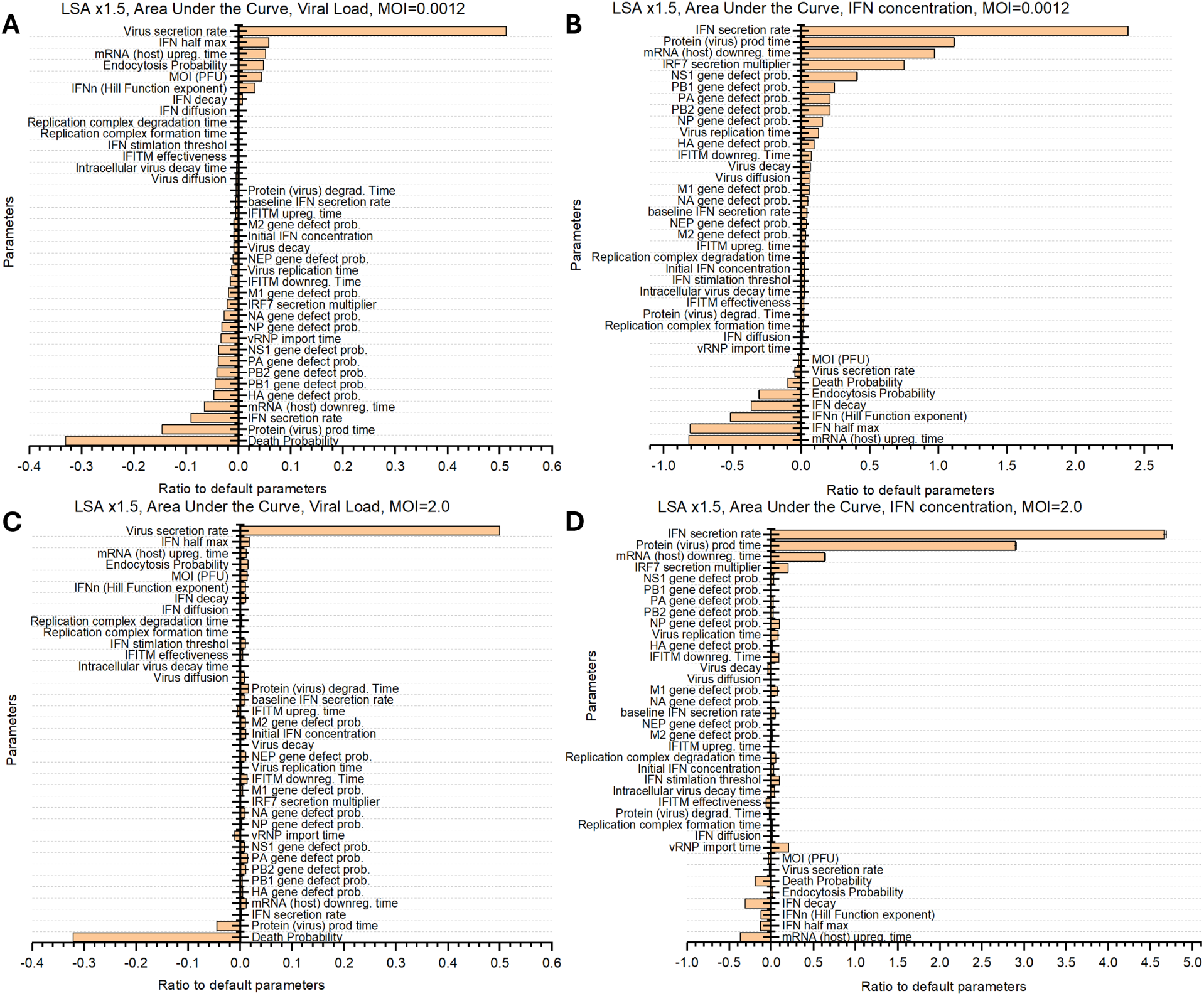
Local Sensitivity analysis for viral load (left column) and IFN concentration (right column) for high (top row) and low (bottom row) MOI. Parameters were individually varied by 50%. Reported metric is the area under the curve (AUC) of viral load or IFN levels, measured as average IFN concentration, relative to default. Averages calculated over 200 replicates for each parameter set. **A, B)** At low MOI, the IFN secretion rate has the biggest impact on the AUC of IFN. **C, D)** At high MOI, the simulation is most sensitive to the IFN secretion rate, average production time of viral proteins, death probability rate, virus secretion rate, and MOI itself.

### Local sensitivity analyses reveal the importance of inoculation MOI

Figure 6 shows the results of a local sensitivity analysis (LSA) on the area under the curve (AUC) of viral load or IFN levels, measured as average IFN concentration, relative to default. At high MOI, the LSA shows that only parameters controlling the IFN secretion rate, virus release rate, cell death probability rate, and the average time of protein and mRNA upregulation significantly affect viral load and IFN levels. In contrast, parameters associated with spatial heterogeneity — such as diffusion coefficients and viral gene defect probabilities — do not significantly affect IFN or viral levels. Because these parameters primarily influence local concentration gradients and spatial variability, their lack of influence indicates that the system dynamics are effectively well-mixed at high MOI. This suggests that the high initial supply of viruses enables intracellular complementation, mitigating the impact of defective genes and reducing the role of spatial structure. Consequently, spatial aspects of the simulation are less important at high MOI than at low MOI.

We expected spatial structure to be more important at low MOI than at high MOI. Our LSA agrees with this prediction, showing a stronger impact of diffusion coefficients and gene defect probabilities on IFN secretion and viral load at lower MOI. Diffusion coefficients are more important at low MOI because virus production and IFN secretion are initially spatially constrained to nearby cells and spread to uninfected tissue sites by diffusion. Gene defect probabilities have more effect on IFN production at low MOI because cells infected by a small number of SIPs are more common, increasing the likelihood that an infected cell does not contain functional NS1. Increasing the viral genome defect rates increases IFN level AUC and decreases viral load AUC at low MOI. The IRF7 multiplier parameter, which increases the rate of IFN secretion, also plays a more important role at low MOI, highlighting that paracrine IFN signaling is intrinsically a spatial process. At low MOI, cells infected with NS1-deficient SIPs have more time to release IFN before complementation shuts off IFN production, and there are larger areas of initially uninfected cells surrounding each infected cell. In these regions, diffusion of IFN can lead to large patches of antiviral-resistant, uninfected cells, making the role of IRF7 in turning on resistance more significant.

The defect probability for the viral gene NS1, responsible for IFN antagonism, has the greatest effect on viral load and IFN production (a higher defect rate leads to less virus and more IFN), followed by those of the genes PB1, PB2, and PA, which are necessary for viral replication. Increasing the rate of death of infected cells (see Supplemental Table 3) and reducing the time during which infected cells produce and release virus represent the most effective ways to reduce viral load AUC. Higher death rates for infected cells also reduce IFN levels, both at high and low MOI. Increasing the cell death rate does not significantly increase the total number of dead cells at the end of the simulation.

Finally, the simulation predicts that increasing the cell’s viral endocytosis probability increases viral load AUC because more rapid endocytosis increases the frequency of superinfection and genome complementation, increasing the likelihood that any given infected cell can release virus (and fail to release IFN), which promotes IAV replication and spread. This result disagrees with other ABMs of viral infection, in which superinfection has no effect on viral replication or IFN production. In these simulations, increasing the endocytosis rate decreases viral load, because faster endocytosis leads to nonproductive superinfection and removes virus from the environment where it could infect other uninfected cells (36,54,60). Our prediction agrees with experiments that study the impact of reducing endocytosis on viral load in cell cultures (69–72), which find a decrease in viral load via the drug chlorpromazine. However, the effect on viral load cannot be exclusively associated with the reduction of endocytosis due to other drug-induced effects (e.g., cytotoxicity).

This LSA underscores the importance of high MOI experiments to decouple parameters that govern individual cell behaviors, such as secretion and cell death rates, from those that influence spatial processes and are, therefore harder to identify, including virus gene defect rates and the diffusion coefficients of virus and IFN. Because spatial effects can be neglected at high MOI, experiments at this regime allow identification of parameters related to individual cell behavior. Performing only high MOI experiments, however, fails to capture spatial effects of IAV infection and IFN response, such as paracrine IFN signaling and SIP-induced IFN production. For that, low MOI experiments are necessary.

## Discussion

Farrell *et al.*’s pioneered IAV infection modeling with SIPs and complementation using a full ODE model (29). They showed that implicit spatial structure (abstracted as a Michaelis-Menten term) is fundamental to IAV spread in the epithelium because complementation of SIPs happens within individual cells. They also discuss the importance of complementation of SIPs for IAV spread. Our simulation, in addition to considering SIPs, also incorporates explicit spatial structure (cells in a lattice) and treats cells as agents. This allows us to explicitly track viral genes within each cell. We additionally include mechanisms for IFN release, viral antagonism, the impact of NS1-defective SIPs on the innate immune response, and passive transport of virus and IFN via diffusion.

Biologically, gene defects can happen for multiple reasons, such as viral ribonucleoprotein (vRNP) complexes improperly coated by nuclear proteins, segment structural damage, truncation, failure to import vRNPs, and other causes that are not necessarily loss-of-function point-mutations due to viral polymerase errors. In all these cases, a single defect in a segment may compromise all genes in the segment, increasing gene defect correlation. Our gene defect model assumes loss-of-function-mutations and neglected defect correlations of genes within the same segment. Future work will need to consider modeling defects that compromise the full segment instead of only the gene. Our simulation allows changing from gene-based defects to segment-based defects easily.

Our results find that MOI plays a critical role in the results of both experiments and simulations for two main reasons. First, the initial phase of plaque growth has an exponential behavior on the viral load; therefore, the time scale of viral load and IFN release is sensitive to the initial fraction of infected cells. Second, different MOI regimes allow identification of different parameters. Higher MOI presents less spatial heterogeneity, helping to decouple parameters associated with spatial effects – e.g. diffusion coefficients – from parameters associated with individual cell behavior – e.g. virus release rates. Low MOI conditions, on the other hand, reveal other aspects of IAV infection, such as the role of SIPs, complementation, diffusion of IFN and virions, and bystander effects of IFN response. Viral infection experiments should explore a wide range of MOI and measure time series for viral load and cytokines for each case. Estimating MOI is challenging, so application of more precise experimental techniques to measure MOI would be helpful.

Unlike previous ABMs (36,54,60), our model predicts that increasing the endocytosis rate (speeding up endocytosis of viruses by epithelial cells) would increase viral load. In ABMs that represent both virus and cells explicitly, higher endocytosis increases superinfection and removes more virions from the environment, reducing the number of virions available to infect other uninfected cells. Therefore, once successfully infected, a cell that keeps endocytosing viruses hinders infection spread. On the other hand, when we consider SIPs, genome complementation becomes essential to productive infection, so superinfection increases virus spread. Experiments are limited, but suggest that decreasing endocytosis decreases viral load (69–72), in agreement with our model.

Unexpectedly, in our experiments, high MOI cultures derived from five different donors had similar IFN levels and viral load curves during IAV infection. To conclude that the differences between donors at the epithelial cell level are negligible, additional samples from a more diverse pool of donors would be necessary. However, sourcing these cells, especially from younger healthier donors, represents a challenge. For future work, we will focus on increasing the sample size of donors, testing low MOI conditions, and exploring other sources of donor heterogeneity, such as innate immune cells (e.g., dendritic cells, macrophages), by including them in the epithelial culture.

In the IFN treatment experiment, viral load at 72 hpi is lower than the viral load predicted by the simulation (Figure). When we run our simulations for long periods with different IFN treatment concentrations, we observe the same viral load saturation value. Our assumptions capture a delay in the onset of virus release but do not capture “subcritical” infection spread (wherein each infected cell infects less than 1 other cell on average) for high doses of IFN treatment. Experiments show that treatment with exogenous IFN-*λ* at high concentrations dramatically reduces viral load (40,73,74), suggesting that IFN treatment could lead to attenuated virus spread. However, there is also evidence that IAV virions can last hundreds of days in epithelium despite innate immune response (75). Sustained infection is likely negated by the presence of an adaptive immune system, which we did not include in the model. Additional experiments measuring plaque arrest outcomes as a function of the concentration of IFN treatment prior to infection are key to improving models of IAV infection. For future work, we plan on incorporating a different set of assumptions that can lead to subcritical infection spread in future simulations.

Our simulation predicts that decreasing the lifespan of infected cells reduces both viral load AUC and IFN AUC. Experimentally, this is supported by the findings of Wong and colleagues that drug candidates that increase stress response to viral infections significantly reduce viral titers in different viral infections (76). Because all infected cells ultimately die in this scenario, such a drug theoretically does not increase tissue damage. Moreover, the decrease in IFN levels could avoid cytokine storms. This intervention strategy could also apply to other types of viruses, as the amount of virus released is associated with the time that the cells spend in the *virus releasing* state. Our approach shows how model predictions could be used clinically or experimentally to prioritize countermeasure development or testing (77).

Representing NS1 antagonism of innate immune response and IFN production driven by NS1-defective SIPs in epithelial cells is essential to the long-term goal of developing a digital twin of IAV infection in cell culture, where the behavior of innate immune cells such as macrophages and neutrophils heavily depends on the release of cytokines and chemokines. In future work, we will experimentally explore differences in innate immune cell response to IAV among different donors in lung epithelium *in vitro* and adapt the model to include immune cells, and modularize our SIP and complementation mechanisms and include them in Sego et al. model (44). Further development of the model will enable valuable biological insights, such as how intracellular competition between defective and complete viral genomes affect viral replication and progeny virion composition; how a compromised gene affects the transcription of other genes in the same segment; and how different types of gene defects impact transcription and viral protein function.

## Acknowledgments

We acknowledge Jason Fair, Erin Conroy, and the Biocomplexity Institute supporting this project.

## Author Contributions

Dal-Castel P.: Conceptualization; Methodology; Software; Validation; Formal Analysis; Investigation; Data Curation; Writing – original draft; Writing – Review & Editing; Visualization

Resnick J.: Conceptualization; Methodology; Validation; Formal Analysis; Investigation; Data Curation; Writing – original draft; Writing – Review & Editing; Visualization

Gallagher M.: Conceptualization; Methodology; Software; Validation; Resources; Data Curation; Writing – original draft; Writing – Review & Editing; Visualization; Supervision; Project administration; Funding acquisition

Sluka J.: Conceptualization; Methodology; Software; Validation; Writing – Review & Editing

Helfers M.: Investigation

Bird I.: Investigation

Grady S.: Conceptualization; Methodology; Validation; Resources; Data Curation; Writing – original draft; Writing – Review & Editing; Visualization; Supervision; Project administration; Funding acquisition

Glazier J.: Conceptualization; Methodology; Software; Validation; Resources; Writing – Review & Editing; Supervision; Formal Analysis; Data Curation

