## Supplemental Materials for "A Simulation of Semi-Infectious Particles and Genome Complementation Reproduces Interferon Response by Respiratory Epithelial Cells *in vitro* during Influenza A Virus Infection"

#### Model Source Code and Movies

Source code for our models, instructions for installing and running, literature, experimental and model derived parameters, representative results and simulation movies are available on Zenodo [https://doi.org/10.5281/zenodo.20303705] and on GitHub [https://github.com/pdalcastel/IAV_Infection_CC3D_MaBoSS_2026].

#### Simulation Movies

Movies are available on Zenodo [https://doi.org/10.5281/zenodo.20303705] and on GitHub [https://github.com/pdalcastel/IAV_Infection_CC3D_MaBoSS_2026].

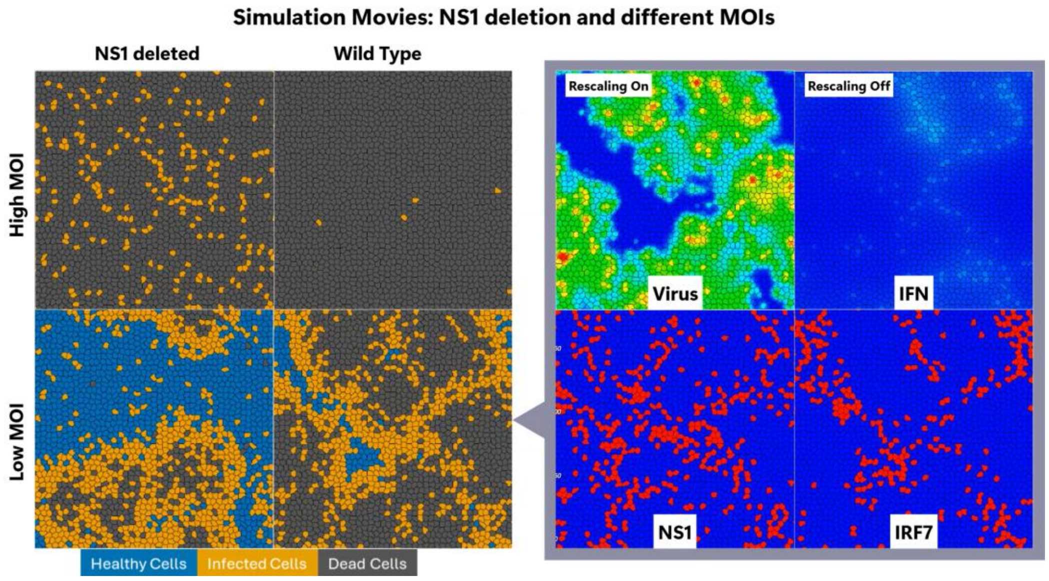


**Supplemental Movie 1.** Simulation movies showing the propagation of wild type or NS1-deleted IAV infection under high/low MOI. NS1 deletion has a significant impact on infection spread, resulting in less virus, fewer dead cells and higher IFN response at both high and low MOI. We observe spatial heterogeneity in IFN response, viral production, and plaque shape. Hotspots of IFN production tend to correlate with regions of the cell culture that only became infected after a delay. At higher MOI, we observe a rapid increase in the dead cell population, with the NS1-deleted infection killing cells in the culture more slowly than WT infection. Our model does not incorporate mechanisms of apoptosis inhibition by NS1.

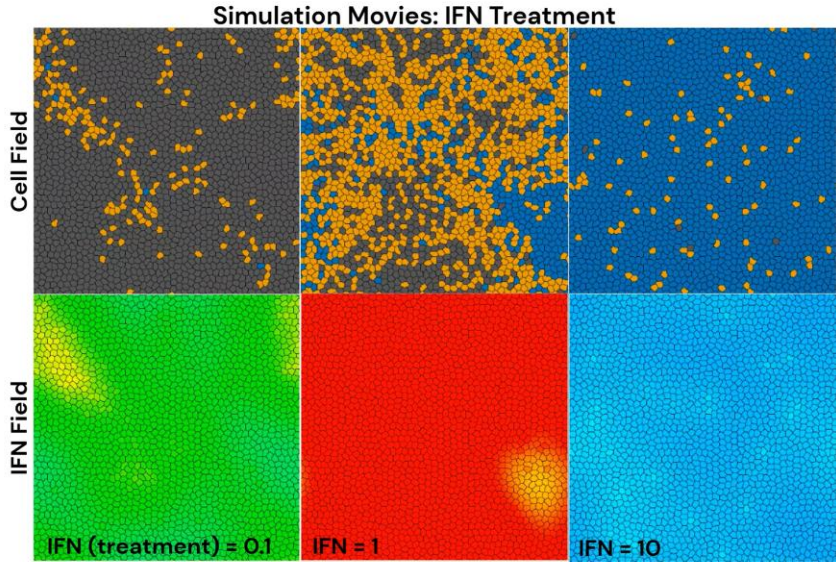

**Supplemental Movie 2.** Intermediate levels of IFN pretreatment caused the largest initial IFN response. The model predicts that high doses of IFN might reduce both extracellular levels of IFN and virus production. However, the model does not consider potential IFN cytotoxicity. The color scale for IFN goes from blue (zero) to red (35 ug/mL).

### Intracellular Pathways

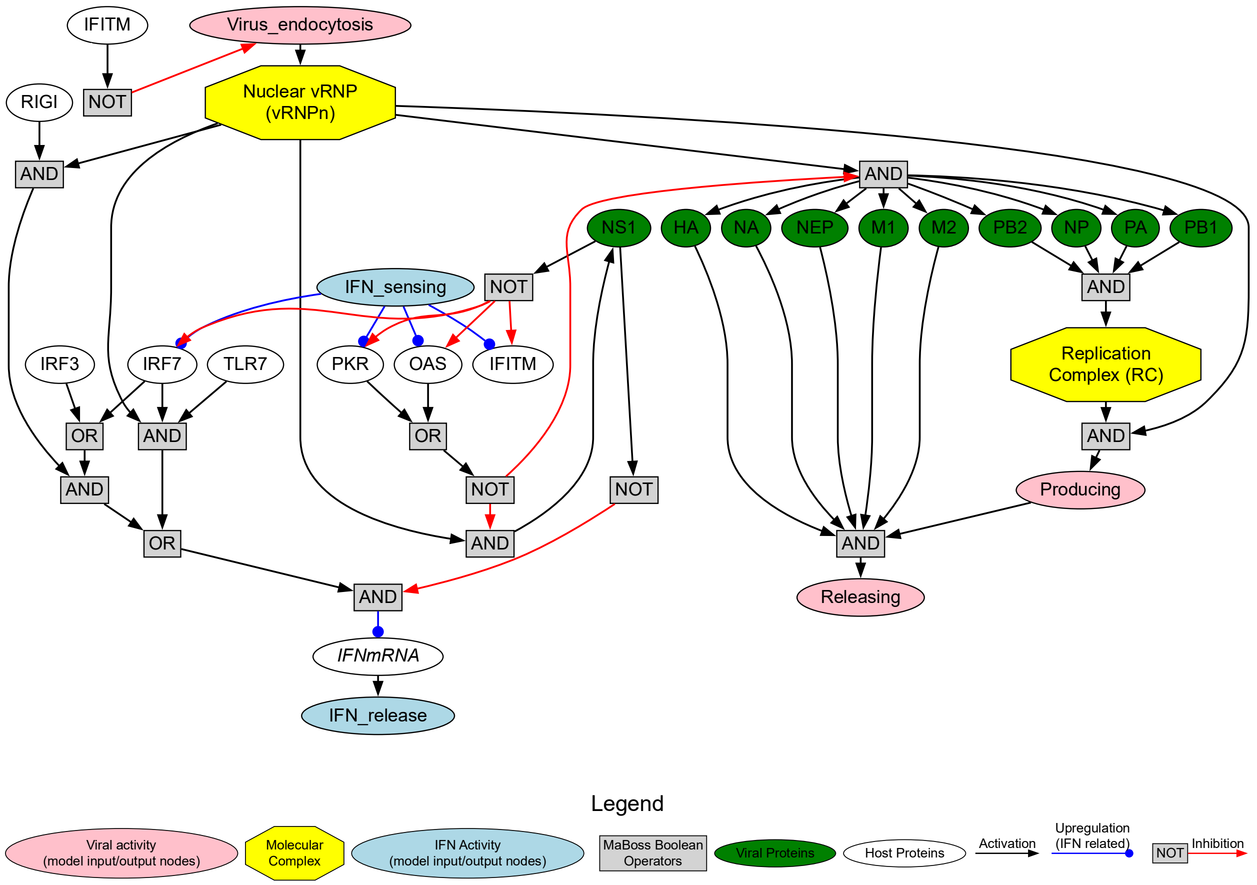


**Supplemental Figure 1**. Boolean stochastic network (**BSN**) diagram of intracellular pathways. This network includes pathways for the virus life cycle, NS1 antagonism, IFN production and IFN-induced activation of ISGs.

### Supplemental Materials Section 1. Gene defect probabilities and complementation:

For IAV, estimates of the ratio of IUs to NIPs vary between 1:10 and 1:1000 (1–3). NIPs have different phenotypic effects on cells, such as inducing or suppressing IFN response (3,4), mainly depending on virus competency in producing NS1 protein (3,5,6). For SARS-CoV-2, Bhat *et al.* estimate 1:10 - 1:100 ratios of IUs to DIPs (7), and report that DIPs decrease viral output. Because of the reduction in viral output, some studies suggest that DIPs could be used to treat infection, but do not specify how and when to deliver the defective viral genomes or DIPs (8,9). Some computational models have implemented DIPs as a separate population of viruses that cannot replicate inside cells on their own, and interfere with the replication of superinfecting infectious virions (10–12). Here, we focus on complementation between different SIPs, so we model SIPs but not DIPs.

Zhaobin *et al.* modeled NIPs of SARS-CoV-2 assuming that genome errors originate from RNA-dependent RNA-polymerase's (RdRp) mutations (13). They coupled their ODE system representing NIP production to a stochastic agent-based epidemiological model for a population of humans and studied how NIPs impact epidemics dynamics. Farrell and colleagues implemented viral complementation of SIPs in a spatially structured ODEs system (14), but assumed that the population of SIPs is homogeneous and did not consider their effect on innate immune response of the host cell. Inspired by Zhaobin *et al.*'s model, we assume a loss-of-function point mutation probability per base and estimate the probability that each gene is defective. Jacobs *et al.* (15) show that a more realistic description would consider segment loss during budding, transport and from degradation by host-cell proteins. We will include segment loss in future work. Regardless of the source of genome errors, cells infected by an NS1-incompetent SIP will produce more IFN than those infected by a wild type virus still holds.

We assume one loss-of-function-mutation for every 1000 bases in the viral genome per replication cycle. From this rate we predict a ratio of IFN suppressing particles per IU of approximately 54:1, and a ratio of IFN inducing particles per IU of approximately 13:1; estimates from experiments are 50:1 and 10-20:1, respectively (1,3). This agreement is enough to capture ratios of IFN suppressing and IFN inducing particles to IU roughly similar to the experimental values during our simulation. Therefore, we accept the assumptions we used to generate these estimates, and we extrapolate them to estimate the defect probability of other viral genomes.

To estimate the gene defect probabilities we assume:

1. The sources of gene defects errors are loss-of-function point mutations,
2. Novel production of RdRp is not required for a SIP to be IFN suppressing or IFN stimulating,
3. The error rate per base is constant,
4. RdRp makes 1 loss-of-function-mutation error for every 1000 bases copied in the viral genome, and
5. NS1 is the main viral protein responsible for inhibiting IFN response

Then we use the equation

$$\mathrm{P(IU)} = (1 - P_{db})^{\mathrm{S}}, \qquad S1$$

**Supplemental Equation 1**

to estimate the probability that a virus an IU. P(IU) is the probability that a virion is an IU, which we obtain from the IU/SIP ratio from the literature; $P_{db}$ is the defect probability per base of the viral genome and S is the total size of the genome.

Next, we use:

$$\mathrm{P(IFN^-)} = (1 - \mathrm{P_{db}})^{S_{NS1}}, \qquad S2$$

**Supplemental Equation 2**

and,

$$\mathrm{P(IFN^-)} = 1 - \mathrm{P(IFN^+)}, \qquad S3$$

**Supplemental Equation 3**

to estimate the probability that a SIP is IFN suppressing ($\mathrm{IFN^-}$) or IFN inducing ($\mathrm{IFN^+}$), where $S_{NS1}$ is the size of the NS1 gene. The defect probability of each gene depends on its size.

Finally, consider that a number $M$ of virions enter the same cell, and each of these virions contains potentially defective genomes. These genomes may or may not complement each other's missing functions. We calculate the probability that $M$ of genomes provide good copies of all $N$ viral genes. Assuming that gene defects are independent, and that genomes are independent, the complementation probability is

$$P(N,M) = \prod_{j=1}^{N} \left(1 - {p_j}^{M}\right). \qquad S4$$

**Supplemental Equation 4.** Complementation Probability.

where $M$ is the number of genomes, $N$ is the number of genes, and $p_j$ is the defect probability of the gene with index $j$. This equation could be used to build simplified versions of our model, without explicit representation of viral genes. In such simplified system, infection related cell phenotypes (IFN producing, NS1 antagonism and virus producing) would depend on the number of infecting genomes, without the need to track every single viral gene present in each cell. We could easily adapt this equation to fit other sources of genome defects such as segment loss. We could also employ multiple instances of this equation, one for each phenotype of interest by organizing the viral genes in broad phenotype categories, such viral antagonism, virus replication and virus release. This equation may be useful to approximate the number of successfully infected cells given an MOI value for high MOI, since the Poisson estimate underestimates the number of infected cells for not considering complementation events.

### Error Threshold Regions

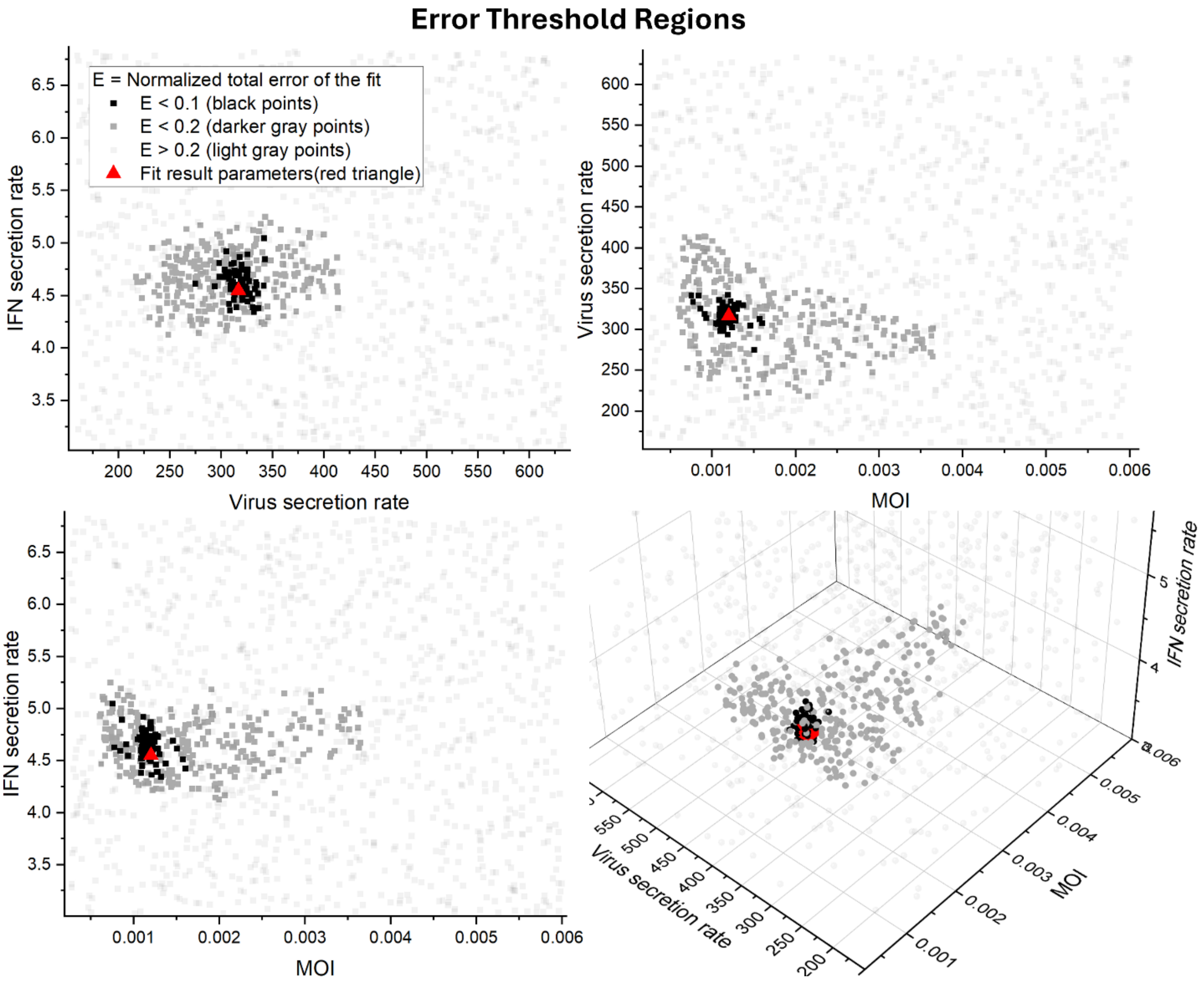


**Supplemental Figure 2.** Error threshold regions illustrate the range of parameter values consistent with the experimental data. Each region indicates the combinations of parameters for which the model's predictions deviate from the experimental measurements by an error value of 0.1, i.e., ±10% (black points) and 0.2, i.e., ±20% (darker gray points). These plots visually summarize parameter uncertainty, correlations between parameters, and whether the fitting parameters fall into a constrained region of the parameter space, helping to identify which parameters are tightly constrained and which are more loosely defined. The error value for each parameter set averages over 10 replicate simulations. The threshold regions shrink as the normalized error tolerance decreases, showing that the fitting parameters fall inside a compact region of the parameter space. The parameters are weakly correlated, and the MOI is the parameter with lowest identifiability.

### Supplemental Table 1. Biological observations and model parameters.

| Viral complementation and Noninfectious Viral Particles | |
|---|---|
| **Observation** | **References** |
| Rate of viral complementation increases with MOI | Marshall N. et al. 2013 (16), Taylor K. et al. 2023 (17) |
| Individual viruses rarely cause productive infections, relying instead on complementation | Phipps K. et al. 2020 (18) |
| Instances of noninfectious viral particles and viral complementation events are frequent | Jacobs N. et al. 2019 (15) |
| Augmented ratio of noninfectious viral particles to PFU is associated with increased inflammatory response in a mouse model | Penn R. et al. 2022 (19) |
| Noninfectious viral particles are diverse in their phenotypic effects, but can be separated into IFN inducing and IFN suppressing categories | Brooke C. 2014 (1) |

| Viral NS1 Protein and IFN Suppression | |
|---|---|
| **Observation** | **References** |
| Intracellular level of NS1 contributes to heterogeneity in single cell response to viral infection | Yang Q. et al. 2023 (20), Russel A. 2019 (21) |
| NS1 is the main factor responsible for suppressing IFN response | Russel A. 2019 (21) |
| Noninfectious viral particles can either suppress or induce IFN. NS1 is necessary to suppress IFN induction | Marcus P. et al. 2005 (3) |
| Infection of IAV with NS1 deletion causes 10-100x more IFN production compared to infection with NS1 viable viruses | Marcus P. et al. 2005 (3), Tawaratsumida K. et al. 2014 (22) |

| | |
|---|---|
| NS1 inhibits IFN production, virus detection, and anti-viral gene expression | Krug R. 2015 (23), Yang Q. et al. 2023 (20) |

| **MOI and IFN response** | |
|---|---|
| **Observation** | **References** |
| Very high and very low MOIs show lower IFN response than intermediate MOIs at 24 hpi | Alalem M. et al. 2023 (24) |
| High MOI gives severe infection with low IFN response, while low MOI gives less severe infection with enhanced IFN response | Grabowski F. et al. 2023 (25), Ramos I. et al. (2019) (26) |

| **IFN Production Mechanisms** | |
|---|---|
| **Observation** | **References** |
| One viral genome is enough to trigger IFN production | Marcus P. 1983 (27), Thoresen D. 2023 (28) |
| IRF7 is necessary for robust and sustained IFN production | Wu W. et al. 2020 (29) |
| Robust IFN production requires detection of the viral genome | Wu W. et al. (2020) (29), Vicary A. et al. (2023) (30) |
| IFN production happens in two phases: 1) virus induced IFN production in initially infected cells, 2) enhanced IFN production by infected cells with high IRF7 and augmented RIGI expression | Wu W. et al. (2020) (29) |

| **ISG Activity and Antiviral Effects** | |
|---|---|
| **Observation** | **References** |

| | |
|---|---|
| ISGs help contain the spread of viruses via: 1) inhibition of virus endocytosis, 2) inhibition of viral gene expression, 3) enhanced IFN production | Wu W. *et al.* (2020) (29), Schoggins J. (2014) (31), Husain M. (2024) (32), He X. *et al.* (2024) (33) |
| Bystander cells show increased ISG activity | Ramos I. *et al.* (2019) (26) |

| **Viral Replication and Production** | |
|---|---|
| **Observation** | **References** |
| Influenza A Virus first synthesizes viral proteins that stabilize the viral genome. Replication of the viral genome starts after protein synthesis | Pflug A. *et al.* 2017 (34), Vreede F. & Brownlee G. 2007 (35) |
| A minority of cells produce most of the virions, and viral production is highly heterogeneous, with viruses produced per cell ranging from 10 to 1000 viruses | Zath G. *et al.* 2024 (36) |

### Supplemental Table 2 Model Assumptions and Simplifications

| **Intracellular Pathways and Host Response** | |
|---|---|
| **Assumption** | **Notes / Limitations** |
| Intracellular pathway nodes that represent proteins and gene expression switch state from inactive to active and active to inactive stochastically and depend on the state of other interacting nodes of the BSN | This simplification of intracellular pathways allows the use of BSNs to model virus-host antagonism. We do not have direct quantitative measurements of intracellular molecules, but we still can reproduce tissue-level IFN and viral load with Boolean representations of protein levels. **Pros:** speed, large number of nodes. **Cons:** unable to model graded effects of key proteins like NS1 and proteins derived from ISGs. Moreover, Stochastic Boolean Networks cannot capture the time delays in transport, transcription and translation. |

| | |
|---|---|
| NS1 is the carrier of IFN inhibition in IAV. When NS1 is active, the host cannot express ISGs, cannot sense viral presence, and cannot express IFN | Other factors may also contribute to inhibition of IFN production, such as primer snatching by RdRp. Different concentrations of NS1 can have different effects and efficacy. |
| Viral genome triggers IFN production. One molecule of viral genome is enough | This assumption simplifies the mechanism of virus detection so that implementation with BSN is straightforward, but does not capture the graded IFN response to different numbers of viral genome inside the cell. |
| ISGs activation rate depends on the local concentration of IFN via a Hill Function | This response mechanism affects the transition probability of ISGs; however, the internal levels of ISGs are still binary. |

| **Viral Population and Representation** | |
|---|---|
| **Assumption** | **Notes / Limitations** |
| The distribution of the frequencies of the different types of SIPs in the population of viruses does not change in time | Experiments suggest that the ratio of noninfectious particles to IU may increase over time and that cells coinfected with noninfectious particles release more noninfectious particles, causing accumulation of NIPs. We neglect this effect because it is mainly a result of DIP action, while our work focuses on SIPs. |
| Representation of viruses as a diffusing concentration field | The continuum approximation introduces abnormalities, such as less than 1 virus in a pixel, which can affect the outcome of a simulation at low MOI. This simplification avoids representing individual virions as agents with random walk dynamics, which is computationally expensive, given the large number of viruses circulating in the simulation (~100k viruses). |
| When a cell takes up a virus, the number of viruses in the field is locally reduced by 1 unit | This assumption is necessary to capture the local decrease in the number of virions in the medium during an endocytosis event. |
| A cell cannot take up virus from the field if the local field amount is less than 1 | This assumption is necessary to avoid negative virus concentrations. |

**Virus Endocytosis and Genome Defects**

| **Assumption** | **Notes / Limitations** |
|---|---|
| Endocytosis is probabilistic with a constant probability rate. When IFITM is active, this rate is lower | This assumption does not capture the time delay of virus internalization and saturation effects, which could affect the dynamics of viral spread. |
| Cells that are releasing virions cannot endocytose additional virions | This assumption comes from the biological hypothesis that cells in advanced stages of infection suppress viral entry, referred to as *superinfection exclusion*, which happens at about 6 hpi (37). |
| We assume that the infection happens mostly on top of the epithelium, and that the mucus layer is very thin | This assumption allows modeling the system in a 2-dimensional lattice, but does not capture effects of the infection spreading to the cells below, and their potential impact on IFN signaling. |
| RdRp point-mutations result in protein loss of function | We assume that either a gene is functional or it is completely non-functional. This makes the problem tractable by allowing us to easily estimate defect probabilities based on available cell phenotype data. We neglect non-fatal mutations, which would partially compromise the function of a protein, impacting the kinetics of reactions in a non-binary "all or nothing" way. |
| When a cell takes up a virus, the presence of productive viral genes is based on an independent draw for the defect probabilities of each gene | This assumption allows us to use a single field to represent the viral population. However, this assumption allows a cell that does not contain the NS1 gene to effectively release viruses that can infect other cells as NS1 competent viruses. All other nine viral proteins are required for the cell to release functional virions. |

**Cell Death and Secretion Dynamics**

| **Assumption** | **Notes / Limitations** |
|---|---|
| A cell that produces virions is susceptible to cell death. The cell | This assumption is based on the biological hypothesis that cells die due to resource reallocation to viral genome replication and viral protein production instead of host metabolism, therefore, we require the cell to be |

| | |
|---|---|
| death probability per unit time is constant | producing virions to be susceptible to dying from the infection. The specifics of how metabolism affects apoptosis during infection are complicated, so we simplify the problem by implementing a constant death probability per unit time. |
| The release rate of virions is constant after the start of virion release | Biologically, the release rate of virions s peaks early, then decreases as the infected cell ages (38). We simplify this process by assuming a constant release rate of virions. |
| Cells do not release virions after death | This assumption is based on the biological hypothesis that IAV budding happens at the cell membrane and is necessary to produce working virions. Non-assembled viral material released from inside a dead cell is not infectious. |
| The secretion rate of IFN is constant and secretion requires that the cell senses the viral genome. With IRF7 present, the secretion rate of IFN increases by a factor of 10 | This assumption is based on the biological hypothesis that IFN production is downstream of RIGI and TLR7 signaling, and that IRF7 boosts IFN production via a positive feedback loop. |

| **Signaling and Sensing** | |
|---|---|
| **Assumption** | **Notes / Limitations** |
| Cells sense the average concentration of IFN locally | We do not explicitly represent membrane receptors for IFN. |
| Dead cells don't take up/secrete diffusive fields and don't participate in signaling pathways | Biologically, dead cells can participate in certain processes such as DAMPs and PAMPs signaling, and they can release material that interferes with the diffusion of viruses. We opted to not include DAMPs and PAMPs since they are not the focus of our work. |
| Cells have a baseline secretion rate of IFN | We assume that all live cells have a constant baseline IFN secretion rate. |

| **Cell Culture and Measurement Simplifications** | |
| --- | --- |
| **Assumption** | **Notes / Limitations** |
| We ignore potential impacts of replacing the culture's medium when a measurement is taken in the *in vitro* experiments | Although replacing the culture's medium may interfere with cell behavior, we do not have information on how it impacts and by how much. To simplify this problem, we reason that replacing the medium removes excess IFN and does not remove IFN that is on the membranes of cells. Therefore, we opted to not explicitly represent wash events in the model. |
| Cell culture is modeled as 2D | In experimental cell culture, cells can stack and the thickness of the cell layer can play a role in virus spread and IFN sensing. We assume that the top cell layer is the most relevant during virus infection and spread, we define a 2D tissue *in silico.* |
| We consider a single cell type culture | Biologically, different cell types have different responses to virus and signals. In our simulation, we consider just epithelial cells that are susceptible to infection. The response heterogeneity of each cell emerges from the stochasticity of the BSN and from the viral genes that are present in the cell. |

### Supplemental Table 3. Model Parameters

| **Genome Defects** | |
| --- | --- |
| **Parameter and Value** | **Reference** |
| NS1 gene defect probability = 0.225 | Estimated from Marcus P. *et al*. 2005 (3) and Brooke C. 2014 (1). |
| PB1 gene defect probability = 0.567 | Extrapolated from the estimated NS1 defect probability and the size of the PB1 gene, see Supplemental Materials Section 1. |
| PB2 gene defect probability = 0.569 | Extrapolated from the estimated NS1 defect probability and the size of the PB2 gene, see Supplemental Materials Section 1. |
| PA gene defect probability = 0.547 | Extrapolated from the estimated NS1 defect probability and the size of the PA gene, see Supplemental Materials Section 1. |

| | |
|---|---|
| NP gene defect probability = 0.425 | Extrapolated from the estimated NS1 defect probability and the size of the NP gene, see Supplemental Materials Section 1. |
| HA gene defect probability = 0.465 | Extrapolated from the estimated NS1 defect probability and the size of the HA gene, see Supplemental Materials Section 1. |
| NA gene defect probability = 0.402 | Extrapolated from the estimated NS1 defect probability and the size of the NA gene, see Supplemental Materials Section 1. |
| M1 gene defect probability = 0.243 | Extrapolated from the estimated NS1 defect probability and the size of the M1 gene, see Supplemental Materials Section 1. |
| M2 gene defect probability = 0.102 | Extrapolated from the estimated NS1 defect probability and the size of the M2 gene, see Supplemental Materials Section 1. |
| NEP gene defect probability = 0.126 | Extrapolated from the estimated NS1 defect probability and the size of the NEP gene, see Supplemental Materials Section 1. |

| **Intracellular Viral Kinetics** | |
|---|---|
| **Parameter and Value** | **Reference** |
| Endocytosis probability = 0.6/h | No references found for this parameter. Considering that around 50% of the virions are internalized after 1h of exposure to the cell membrane (39), and considering the mucus barrier, we choose a pessimistic endocytosis probability of less than 1 virus per hour. |
| vRNP average import time = 3h | Ramos I. et al. (2019) (26), Baccam P. et al. (2006) (40). |
| After vRNP import, avgerage time to viral replication = 1h | Plausible physiological value. Heldt F. et al. (2012) (38), Baccam P. et al. (2006) (40). |
| Intracellular virus average. degradation time = 10h | Plausible physiological value (38). |
| Viral protein average degradation time = 24h | Plausible physiological value (38). |

| | |
|---|---|
| Replication complex average formation time = instantaneous | Binding between these three proteins is much faster than transcription and translation. |
| Replication complex average degradation time = 10h | Plausible physiological value (38). |
| Virus release rate = 10-1000 (literature informed) [PFU per cell per hour], 1900 (fit result) [viruses of any type per cell per hour] | Zath G. *et al*. 2024 (36), Baccam P. *et al*. (2006) (40). We use these values as initial guesses, and we later fit this parameter to our experimental data. |

| **IFN Production and Response** | |
|---|---|
| **Parameter and Value** | **Reference** |
| IFITM effectiveness = 96% | Meischel T. *et al*. 2021 (41). |
| IFITM average upregulation time = 12h | Meischel T. et al. 2021 (41). |
| Host mRNA average upregulation time = 4h | Plausible physiological value, Cheng Z. et al. (2016) (42), Hao S. & Baltimore D (2009) (43), Yang E. (2003) (44). |
| IFITM average downregulation time = 24h | Meischel T. et al. 2021 (41). |
| Host mRNA average degradation time = 12h | Plausible physiological value, Hao S. & Baltimore D (2009) (43), Yang E. (2003) (44). |
| IFN secretion rate = 16 (calibrated from literature data), 22.8 (fit result) [µg/(mLh)] | Estimated from Wu W. et al. 2020 (29), Alalem M. *et al*. 2023 (24), Yang Q. *et al*. 2023 (20). We match the ratios of peak IFN concentration during infection and the half maximum interferon concentration for activation of ISGs. |
| IRF7 boosts IFN secretion rate by a factor of 10 | Wu W. *et al*. 2020 (29). |

| | |
|---|---|
| Baseline secretion of IFN = 0.05 (literature informed), 0.01 (from our experiments) [μg/mL] | Wu W. *et al.* 2020 (29). This is the rest value of IFN concentration in the steady state in the absence of infection. |
| Half maximum IFN concentration = 1 [μg/mL] | This is a fixed parameter. We adjust the IFN secretion rate such that the ratio between the peak IFN concentration to the IFN half maximum in the simulation matches the literature. We acquire the units of [μg/mL] for the Half maximum IFN concentration parameter after comparing the ratios. |
| IFNn = 4 | The exponent of the IFN hill function for ISG activation. No references found for this parameter. |

| **Extracellular Environment and Cell Death** | |
|---|---|
| **Parameter and Value** | **Reference** |
| IFN diffusion = 31 [pixels^2/MCS]. Equivalent to 300 [um^2/h] estimated from hydrodynamic radius and converted to simulation units | Extrapolated from experimental value of virus diffusion using Stokes-Einstein relation |
| Virus diffusion = 0.63 [pixels^2/MCS]. Equivalent to 6 [um^2/h] estimated from Mean Square Displacement of influenza viral particles in mucus | Kaler L. *et al.* 2022 (45) |
| IFN decay = 1/(54h) estimated from IFN time-course curve | Marcus P. *et al.* 2005 (3) |
| Virus decay = 0 | We assume that virus decay is negligible over the time scale of the simulation |
| MOI (considering only PFU) = 0.0012 | This is a free parameter, i.e., we obtain it by fitting the simulation to time series of viral load and IFN levels |
| Cell death probability for infected cells = 0.2/h | Estimated from Serrano J. *et al.* 2021 (46) and Baccam P. *et al.* (2006) (40) |

| **Initial Conditions** | |
|---|---|
| **Initial Conditions** | **Value** |
| Initial states of network nodes related to viral genome, viral genes and viral proteins | 0 |
| Initial states of host cell network nodes RIGI, IRF3 and TLR7 | 1 |
| Initial virus field local amount | 0 |
| Initial IFN concentration field | Equal to the basal secretion rate of IFN divided by the IFN decay rate |
| Cell configuration | All uninfected naive (U) cells start out as 7x7pixel squares |

| **Events** | |
|---|---|
| **Events** | **Description** |
| Infection starts after a relaxation period of 50 MCS, equivalent to around 8h | The simulation randomly seeds viruses on the lattice pixels. The total number of seeded viruses is the product of the total number of cells and the total MOI, which combines PFUs and SIPs. The first time a virus is seeded on a pixel, the seeded amount is 1.99 to avoid the loss of virions to diffusion (a cell can only endocytose integer values of virus). The initial relaxation stage of the simulation allows the cells to relax to relatively hexagonal shapes and the internal cell states to relax. |
| Until MCS=60, the endocytosis probability is 1. At MCS=60, the endocytosis probability reverts to its set value | This change in probability avoids artifacts introduced by representing viruses as diffusive fields, such as losing virions to diffusion diffusion (a cell can only endocytose integer values of virus). The concentration field allows virus level between zero and 1, and at low MOI, the infection results from a small number of virions. We set a high endocytosis probability initially to avoid a wide variation in the initial number of infected cells, and therefore getting more consistent results. |

| **Other Configurations/Parameters** | |
|---|---|
| **Other configurations/parameters** | **Value** |
| Lattice size | 301x301 pixels (90,601 pixels$^2$) – 1 pixel is equivalent to 1.26 microns$^2$ |
| Cell volume constraint | Target Volume = 50, Lambda Volume = 10 |
| Contact energies | J=10 between type “cell” and type “medium”, J=20 between type “cell” and itself |
| Contact energy neighbor order | 2 |
| Cell types | Medium, U (uninfected naive), I (infected), E (uninfected exposed to interferon), D (dead). In practice, U and E are both considered healthy cells, so we use the same color and denomination “H” for both |
| Potts neighbor order | 1 |
| Potts “temperature” (intrinsic cell motility parameter) | 10 |
| Diffusion solver | CC3D’s DiffusionSolverFE |
| Diffusive field boundary conditions | Periodic in X and Y |
| Cell lattice boundary conditions | Periodic in X and Y |
